# Argonaute proteins regulate the timing of the spermatogenic transcriptional program

**DOI:** 10.1101/2024.12.31.630913

**Authors:** Maria de las Mercedes Carro, Alexis Dziubek, Amanda Touey, Elizabeth A. Popkowski, Mark Abdelmassih, Leah Simon, Stephanie Tanis, Faraz Ahmed, Jen K. Grenier, Andrew Grimson, Paula E. Cohen

## Abstract

Argonaute proteins are best known for their role in microRNA-mediated post-transcriptional gene silencing. Here, we show that AGO3 and AGO4, but not AGO2, localize to the sex chromatin of pachytene spermatocytes where they are required for transcriptional silencing of XY-linked genes, known as Meiotic Sex Chromosome Inactivation (MSCI). Using an *Ago413^-/-^* mouse, we show that AGO3 and AGO4 are key regulators of spermatogenesis, orchestrating expression of meiosis-related genes during prophase I while maintaining silencing of spermiogenesis genes. Premature overexpression of spermiogenesis genes during prophase I in *Ago413^-/-^* mice results in subfertility, altered sperm morphology and reduced fertilization capability. We also identify BRG1, a BAF complex subunit, as an AGO3 interactor. Loss of AGO3 and AGO4 results in increased BRG1 in spermatocytes, suggesting that AGO3 aids in removing BRG1 from the XY chromatin to achieve MSCI and demonstrating a meiotic role for AGO3 in transcriptional control through the chromatin remodeling machinery.

## INTRODUCTION

Mammalian spermatozoa are male gametes that are critical for sexual reproduction. To progress correctly through spermatogenesis, a series of distinct and stringently regulated developmental programs must be executed in a precise spatiotemporal manner. Spermatogenesis is divided into three phases: first, proliferation and differentiation of spermatogonial stem cells to produce primary spermatocytes. Next, primary spermatocytes enter meiosis, during which they replicate their DNA and undergo two consecutive divisions to form haploid round spermatids. In the third phase, spermiogenesis proceeds as a series of morphological and physiological changes in which round spermatids become elongated spermatids before generating functional spermatozoa ^1^.

Meiosis represents one of the most specialized stages of spermatogenesis with respect to changes in gene regulation and cell biology. Prophase of the first meiotic division requires a series of unique nuclear events to achieve accurate homologous chromosomal segregation during metaphase I. During leptonema and zygonema of prophase I, homologous chromosomes pair and undergo physical tethering, or synapsis, which is achieved through the assembly of a tri-partite protein structure called the synaptonemal complex (SC). Concurrent with SC assembly is the stepwise recombination process that leads to reciprocal exchange of DNA via crossover formation ^2^. Together, SC formation and crossovers serve to maintain homolog interactions through pachynema and diplonema, resulting in the appearance of physical chiasmata structures by diakinesis ^3^.

In male mammals, differences in size and sequence of the sex chromosomes limit synapsis to a small region of homology, referred to as the pseudo autosomal region ^4^. The remaining portions of the X and Y remain unsynapsed, triggering formation of a heterochromatic domain, the sex body, which restricts transcriptional activity of the sex chromosomes ^5,6^. This process, called Meiotic Sex Chromosome Inactivation (MSCI), is initiated by a feed-forward mechanism led by proteins of the DNA Damage Repair pathway (DDR) ^7–9^. Initially, DDR proteins are recruited to the sex chromosome axes where histone H2AX is phosphorylated, generating γH2AX ^10^. The DDR signal spreads to chromatin loops along with additional repressive chromatin marks, establishing the transcriptionally silenced sex body. Following meiosis, XY silencing is partially maintained, resulting in a repressed state of transcription associated with persistence of repressive histone marks during spermiogenesis ^11,12^. The importance of proper establishment of MSCI is highlighted by the existence of checkpoints that cause cell cycle arrest and elimination of germ cells when MSCI defects arise, resulting in infertility ^7,13–16^. Although we now better understand the mechanism of the DDR cascade that leads to MSCI, the epigenetic mechanisms that underlie the chromosome-wide gene silencing during MSCI and maintain low transcriptional activity throughout and beyond pachynema have not yet been fully elucidated.

Previous work has shown that the Argonaute (AGO) proteins, AGO3 and AGO4, are localized to the nucleus of pachytene spermatocytes, where they accumulate in the sex body ^17^. AGO proteins are RNA binding proteins with high similarity, and paralogs in mammals exhibit diverse expression patterns ^18^, suggesting a degree of functional specialization. AGOs are best known for their role as partner proteins with microRNAs, mediating post-transcriptional gene silencing in the cytoplasm ^19–21^. In addition to their participation in post-transcriptional regulation, AGO proteins play a role in transcriptional regulation. In *S. pombe*, AGO-dependent small interfering RNA (siRNA) mediated silencing is required for heterochromatin assembly at centromeric repeats ^22,23^. Furthermore, in *C. elegans*, Argonaute proteins direct the nuclear RNAi machinery to nascent transcripts to promote co-transcriptional repression and recruit histone modifiers ^24–26^. The nuclear role of AGOs in mammalian cells, including in the germline, is less understood. Recent work has suggested AGO involvement in both enhancing and repressing transcription ^27–29^, but the underlying mechanisms and the extent to which mammalian AGOs possess non-canonical nuclear roles has not been established. Reports of germline nuclear localization of AGO2 ^30^ as well as AGO3 and AGO4 ^17^ imply a nuclear role that may be specific to germ cells. However, the lack of reliable tools to study AGO proteins *in vivo* has limited our knowledge of how these proteins contribute to the unique modes of gene regulation found in the germline.

We previously generated mice harboring an *Ago4* deletion, which allowed us to identify a role for AGO4 in MSCI ^17^. In the absence of AGO4, *Ago3* expression was upregulated exclusively in testicular germ cells, along with a concomitant rise in the abundance of AGO3 in the sex body of prophase I spermatocytes, suggesting that AGO3 compensates for loss of AGO4 ^17^. To further investigate the role of AGO3 and AGO4 in sex chromosome silencing, here we examine a mouse strain bearing a deletion of chromosome 4 encompassing the loci encoding *Ago4, Ago1* and *Ago3 (Ago413^-/-^)*. Importantly, previous data indicate that *Ago1* is not expressed in the germline from the spermatogonia stage onwards ^31^, allowing us to use this model to study the roles of *Ago4* and *Ago3* in spermatogenesis. Using cytogenetic, histological, transcriptomic and proteomic approaches, we show that AGO3 and AGO4 are present in the nucleus of prophase I spermatocytes, and that loss of both proteins leads to dysregulation of the gene regulatory program that drives spermatogenic differentiation. AGO3 and AGO4 localize to the sex chromosomes of pachytene and diplotene spermatocytes and their dual absence greatly accentuates the MSCI defects observed in single *Ago4* knockouts. Profiling the interactome of AGO3 and AGO4, along with analysis of regulatory regions of differentially expressed genes in *Ago413^-/-^* germ cells, identifies a nuclear role for AGO3 involving an interaction with the chromatin remodeling machinery through BRG1, a component of the BAF chromatin remodeling complex. Taken together, these data lead us to propose a model whereby AGO3 and AGO4 play regulatory roles at the transcriptional level during meiosis in addition to their canonical roles in post-transcriptional regulation.

## RESULTS

### Argonaute protein localization in germ cells

To investigate the role of AGOs in the male germline, we used a triple knockout mouse strain for the *Ago4*, *Ago1* and *Ago3* genes (Strain # JAX:014152, MGI:101977), which are found in tandem on chromosome 4 (Figure S1A). Deletion by Cre-mediated recombination within exon 6 of *Ago3* and exon 15 of *Ago4* fully removed the intervening *Ago1* locus and the expected portions of *Ago3* and *Ago4* (Figure S1A). Using antibodies specific to each AGO protein, we performed western blot analysis of whole testis lysates and confirmed the loss of AGO3 and AGO4 in *Ago413^-/-^* males (Figure S1B-D). However, we found these antibodies unsuited to immunofluorescence localization in male germ cells. Thus, we generated an *Ago3* tagged mouse line by N-terminal insertion of *myc* and *flag* coding sequences into *Ago3* (*Ago3*^myc-flag/myc-flag^). Additionally, we utilized a previously characterized *Ago2* HA epitope-tagged mouse line (*Ago2*^ha/ha^) ^28^. Notably, western blot analysis of germ cells from *Ago3*^myc-flag /myc-flag^ and *Ago2^ha/ha^* males revealed the presence of AGO3 and AGO2 in both nuclear and cytoplasmic fractions (Figure S2A, B).

To explore AGO3 localization in the spermatocyte nucleus, we performed prophase I chromosome spreads, probing with an anti-FLAG antibody in *Ago3*^myc-flag/ myc-flag^ males in combination with an anti-SYCP3 antibody. We observe no AGO3 in the nucleus of leptotene and zygotene spermatocytes (Figure 1A), and nuclear staining only becomes apparent in early pachynema, when AGO3 associates with the SCs of both autosomes and sex chromosomes. AGO3 localization to the X and Y chromosomes, as assessed by anti-FLAG staining, persists throughout pachynema, spreading through the sex body domain by mid and late pachynema, but is largely absent by diplonema. In contrast, staining of testis sections and chromosome spreads from the *Ago2^ha/ha^* mouse line shows that AGO2 has mainly cytoplasmic localization in the germline, with diffuse nuclear signal seen in prophase I spreads (Figure S2 C, D).

**Figure 1.**
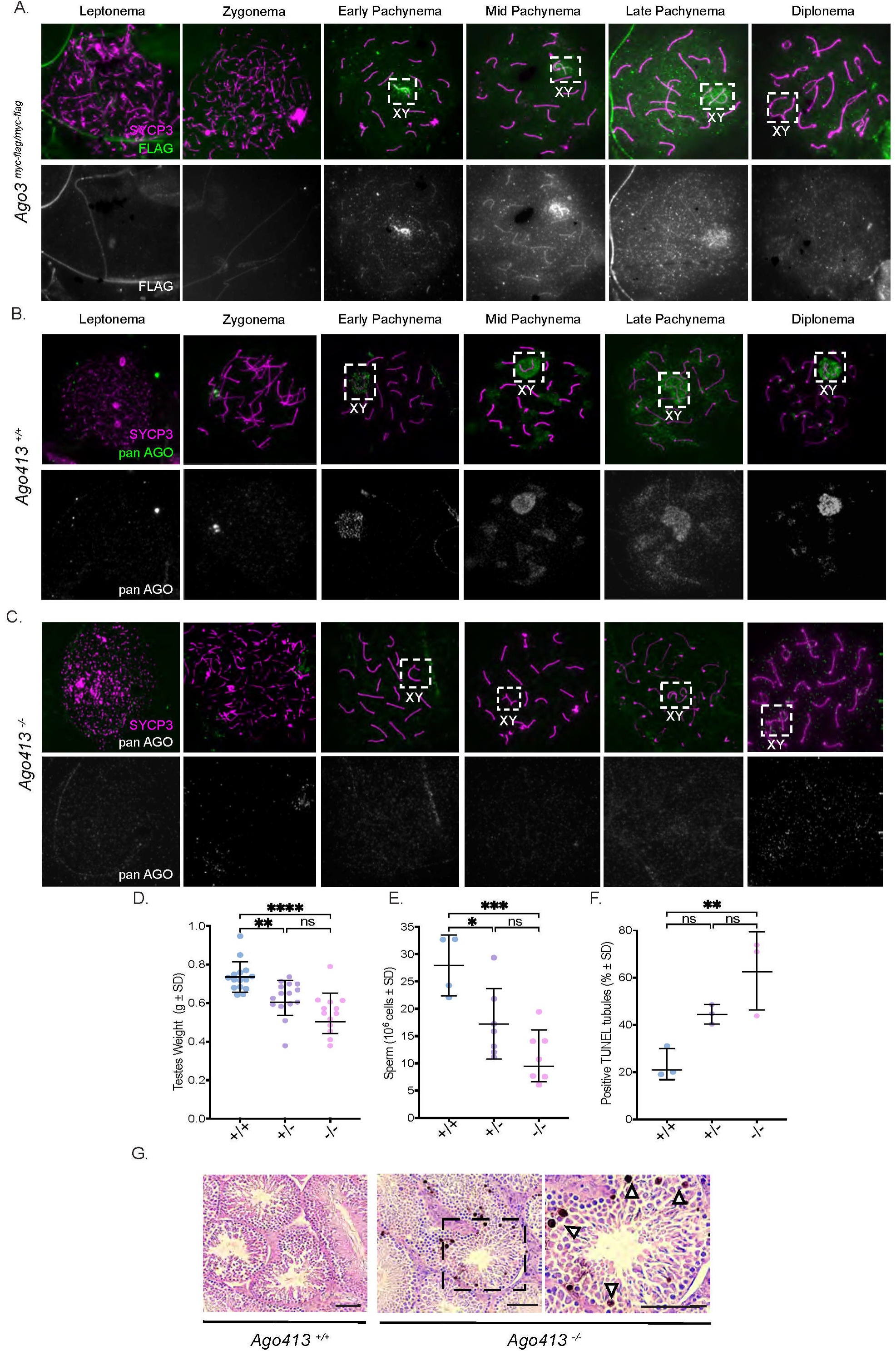
AGO localization during spermatogenesis and *Ago413^-/-^* subfertility phenotype. (A) Meiotic spreads of *Ago3^myc-flag^* mice immunostained with anti-SYCP3 and anti-FLAG antibodies, showing AGO3 localization in indicated substages of Prophase I. (B, C) Meiotic spreads of *Ago413^+/+^* (B) and *Ago413^-/-^* (C) mice; otherwise as described in panel A. (D) Testes weights relative to body weight for indicated genotypes. (E) Epididymal spermatozoa counts obtained by swim out for each genotype. (F) Percentage of tubules showing at least one positive TUNEL cell per each genotype. Dots represent individual measurements and bars represent the mean ± SD. Data were analyzed by ANOVA with Tukey adjustment (**p <0.01, ****p <0.0001). (G) TUNEL staining, bars indicate 40 μm. Insert shows magnification of the *Ago413^-/-^* panel; arrows indicate apoptotic cells.

To further characterize the localization of all four AGO proteins by immunofluorescence, we used a pan-AGO antibody, which confirms AGO localization in the nucleus of prophase I spermatocytes by early pachynema, with this signal persisting through diplonema (Figure 1B). In addition, pan-AGO staining is present on the pericentromeric heterochromatin in mid-late pachytene to diplotene stage spermatocytes. In spermatocytes from *Ago413^-/-^* males, we observe loss of the XY and pericentromeric heterochromatin pan-AGO signal (Figure 1C). Given that *Ago1* is not detectable in wild-type spermatocytes ^31,32^, we attribute the signal observed with the pan-AGO antibody within heterochromatin to AGO3 and/or AGO4, and not AGO1 or AGO2. Taken together, we conclude that AGO3 localizes to the SC of all chromosomes and the sex body during pachynema, while AGO4 localizes to the sex body and the pericentromeric heterochromatin in both pachytene and diplotene spermatocytes. Interestingly, AGO3 alone shows nuclear localization in post-meiotic cells, exhibiting a pattern of focal localization near the chromocenter of round spermatids (Figure S2 E).

### Reduced testis weights and epididymal spermatozoa counts in Ago413^-/-^ mice

In mammals, germ cells mature within the testicular seminiferous tubules, which provide an organized three-dimensional structure that includes germ cells at different stages of development. *Ago413^+/-^* and *Ago413^-/-^* adult mice show decreased testes weight compared to *Ago413^+/+^* males (p <0.01 and p <0.0001, respectively) and have lower sperm counts obtained by extrusion from the cauda epididymis (p <0.05 and p <0.001, respectively; Figure 1 D, E). Although loss of *Ago4* alone causes a subfertility phenotype, including reduced testes weights and epididymal sperm counts ^17^, *Ago413^-/-^* males exhibit a more pronounced phenotype, with an almost 30% reduction in testes weight (Figure 1D), compared to only a 13% reduction in *Ago4*^-/-^ males ^17^. Similarly, sperm counts of *Ago41*3^-/-^ males are reduced by 60% compared to wild-type (Figure 1E), while *Ago4*^-/-^ mice show a more modest 22% reduction ^17^. Histological sections of *Ago413^+/-^* and *Ago413^-/-^* testes reveal no differences in cellular architecture of the seminiferous tubules when compared to wild-type males (Figure S3). However, TUNEL analysis reveals an increase in apoptotic cells in the prophase I to metaphase I layer of the seminiferous tubules of *Ago413^-/-^* mice compared to wild-type males (p <0.01, Figure 1F). This layer is comprised of the cells between the spermatogonia cell layer near the basal membrane and the round spermatid layer near the tubule lumen (Figure 1G).

### Gene dysregulation in Ago413^-/-^ germ cells

To evaluate the impact of AGO3/4 loss on the germline transcriptome, we performed single cell RNA sequencing (scRNA-seq) of enriched germ cell suspensions from testes of *Ago413*^-/-^ and *Ago413*^+/+^ adult males (Figure 2A). We then classified cell clusters into different germ cell populations based on marker gene expression ^33^ (Figure S4A). As expected, *Ago1* expression in wild-type males is confined to spermatogonia while *Ago3* and *Ago4* are detected in spermatocytes from leptonema to pachynema (Figure 2B). We analyzed cell-type composition changes in both the scRNA-seq data (Figure S4B) and in chromosome spreads stained with antibodies against SYCP3 (Figure 2C). Immunofluorescent staining of prophase I chromosome spreads reveals a decrease in the diplotene cell population in *Ago413^-/-^* males (p <0.0001; and also in heterozygotes, p <0.001) compared to wild-type littermates, with a concomitant increase in the pachytene cell population (p <0.05). This change in germ cell proportions during meiosis indicates that *Ago413*^-/-^ spermatocytes are progressing slower from pachynema into diplonema and/or are dying after pachynema.

**Figure 2.**
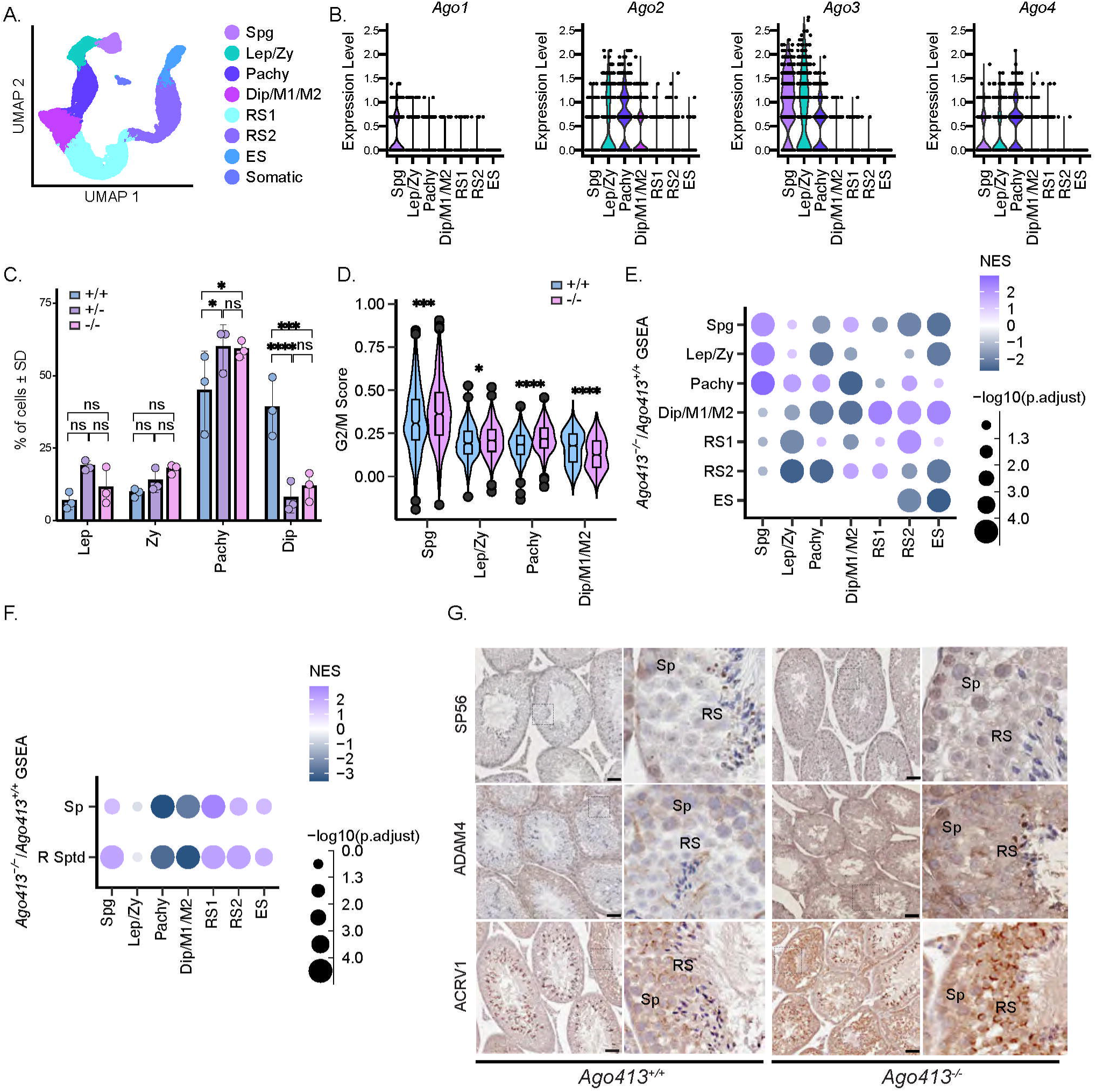
Single Cell RNA-seq analysis of *Ago413^+/+^* and *Ago413^-/-^* germ cells. (A) Uniform Manifold Approximation and Projection (UMAP) plot of cell type groupings in scRNA-seq. (B) Expression of Argonaute genes in *Ago413^+/+^* germ cell types. (C) Percentage of cells in each substage of prophase I obtained after meiotic scoring of prophase I spreads through SYCP3 immunostaining. Bars represent the mean and dots each value ± SD, n=3. Data were analyzed by ANOVA followed by multiple comparisons (*p <0.05, ***p <0.001). (D) Violin plot of G2/M phase marker gene set scores separated by genotype. Bonferroni-adjusted p-values obtained using Wilcoxon rank-sum test (*p <0.05, ***p <0.001, ****p <0.0001). (E) Dot plot showing normalized enrichment score (NES) of cell type marker gene set enrichment analysis. Marker gene sets on X axis and cell clusters on Y axis. Gene sets obtained from wild-type cell marker gene testing (n= for Spg, Lep/Zy, Pachy, Dip/M1/M2, RS1, RS2, and ES respectively). P-values are FDR adjusted and represented by circle size. (F) As in (E) but using bulk RNA-seq lof2 fold-change for testing. (G) Immunohistochemistry staining of testicular sections from *Ago413^+/+^* and *Ago413^-/-^* males using antibodies against spermiogenesis proteins which genes are upregulated in the scRNA-seq and bulk RNA-seq data in the *Ago413^-/-^* pachytene spermatocyte fraction: *Sp56*, *Adam4*, and *Acrv1*. Bars indicate 40 μm, magnified panel of each image show the spermatocytes (Sp) and round spermatid (RS) layers of the seminiferous epithelium where increased IHC signal is observed in *Ago413^-/-^* males.

Using our scRNA-seq data, we analyzed cell cycle progression across the different cell clusters by scoring cells for G2/M cell cycle marker genes. We identified significant differences in the G2/M score in germ cells from *Ago413*^-/-^ males compared to wild-type cells in clusters which are in G2/M or meiotic-equivalent cell phases (Figure 2D). We note a significant increase in expression of G2/M markers in the pachytene cell cluster, indicating higher levels of cell cycle transcripts. We also observe an increase in G2/M marker expression in *Ago413*^-/-^ spermatogonia, which could be due to changes in mitosis between *Ago413*^-/-^ and *Ago413*^+/+^ samples in this cell cluster. In contrast, G2/M markers show a decrease in expression in diplotene spermatocytes, indicating a premature downregulation of cell division-related genes, reflecting dysregulated progression through M phase. We also see that expression levels of specific genes known to be important for spermatogonial differentiation and meiosis, such as Hmgb2 and Top2a, are correlated with marker score (Figure S4C) and are therefore relevant to changes in spermatogenesis. G2/M marker score comparisons overall are consistent with altered progression through the spermatogonia stage and meiosis in *Ago^-/-^* males compared to wild-type.

To investigate how alterations in cell type proportions and cell cycle marker expression levels are related to changes in expression of gene regulatory programs in *Ago413*^-/-^ males, we first identified the top non-overlapping upregulated genes for *Ago413 ^+/+^* germ cells in each cluster, compared to gene expression in other clusters, to identify marker gene sets defining wild-type expression in each cluster (Table S1). We then performed gene set enrichment analysis (GSEA) to test if these marker gene sets showed enrichment in differentially upregulated or downregulated genes in *Ago413^-/-^* cells compared to *Ago413 ^+/+^* cells for a given cluster, indicating abnormal expression of gene regulatory programs (Figure 2E). Consistent with slower progression through earlier stages of prophase I, genes upregulated in leptotene/zygotene and pachytene spermatocytes from *Ago413^-/-^* mice compared to *Ago413^+/+^* mice are enriched for spermatogonial marker genes, while genes upregulated in the *Ago413^-/-^* pachytene cluster also show enrichment of leptotene/zygotene stage markers. Thus, in these earlier stages of meiosis, germ cells in *Ago413^-/-^* animals do not fully downregulate genes which in wild-type animals are normally preferentially expressed in earlier stages of meiosis and become repressed as meiosis proceeds. In contrast, spermatocytes at the diplotene stage through to metaphase II (M2) from *Ago413^-/-^* males display premature upregulation of programs in cells undergoing spermiogenesis. This premature upregulation of spermiogenesis genes in the *Ago413^-/-^* germline continues through to early spermatid development, with earlier round spermatids exhibiting upregulation of late round spermatid markers. Taken together, these results indicate that the normal timing of gene expression transitions in the germline is perturbed in Ago*413^-/-^* animals.

We validated these aberrant gene expression signatures using RNA-seq of pachytene/diplotene spermatocytes and in round spermatids enriched using a densitometry BSA gradient ^34^. We then calculated gene expression differences between Ago*413^-/-^* and *Ago413^+/+^* spermatocytes and Ago*413^-/-^* and *Ago413^+/+^* round spermatids and performed GSEA using the same marker gene sets identified using our scRNA-seq data (Figure 2F). In accordance with our scRNA-seq data, upregulated genes in both Ago*413^-/-^* spermatocytes and spermatids are enriched in marker genes characteristic of wild-type spermatids, indicating premature expression of genes characteristic of later germ cell development stages. We also performed gene ontology term GSEA and observe enrichment of terms in genes upregulated or downregulated in *Ago413^-^*^/-^ males compared to wild-type males (Table S2). In the spermatogonia, leptotene/zygotene, and pachytene clusters we see depletion of terms associated with spermatid development, which may be indicative of delayed transcription that is normally transcribed early in spermatogenesis and stored for later translation (Figure S5A, B, C). The diplotene to M2 cluster shows enrichment of gene ontology terms associated with spermiogenesis (Figure S5D), particularly upregulation of genes needed for sperm flagella development, motility and fertilization processes, consistent with the early upregulation of round spermatid marker genes, since many cellular changes related to proper sperm function take place at the round spermatid stage. The round spermatid stages show enrichment of upregulation in translation and RNA processing (Figure S5 E, F), which are important for regulation during the round and elongating spermatid stages when transcription is reduced. To validate changes in transcriptome profiles of spermiogenesis genes, we performed immunohistochemistry of ADAM4, which is involved in sperm migration and binding to the zona pellucida in the oocyte ^35^, and the acrosomal proteins ACRV1 and SP56 ^36,37, 35^ (Figure 2G). Increased IHC signal for all these proteins in *Ago413^-^*^/-^ testes sections confirm early upregulation of proteins associated with spermiogenesis in *Ago314^-/-^* germ cells.

Overall, analysis of cell cycle markers indicates that spermatocytes from *Ago413*^-/-^ males have a slower progression during the first stages of prophase I into pachynema, which then transitions to an altered expression profile in diplotene spermatocytes in which spermiogenesis genes are mis- expressed during meiosis. These data, plus the existence of checkpoints at the pachytene stage that trigger apoptotic pathways in defective cells ^38,39^, suggest that the increase in cell death detected by TUNEL in the prophase/metaphase layer of the seminiferous tubules of *Ago413*^-/-^ males corresponds to cells failing to transition from pachynema to diplonema.

### Impaired MSCI in Ago*413^-/-^* mice

To progress to diplonema, spermatocytes must achieve full synapsis along all autosomes and within the pseudo autosomal region of the sex chromosomes by the end of pachynema ^13,40,41^. The alteration in prophase I progression of *Ago413^-/-^* males, AGO3 and AGO4 localization to the pachytene nucleus, and the disrupted sex chromosome silencing observed in Ago*4^-/-^* males ^17^, led us to investigate XY-linked gene expression and MSCI during spermatogenesis in *Ago413^-/-^* males. To examine MSCI in *Ago413^-/-^* males, we analyzed XY-linked gene expression in our scRNA-seq data. As we previously observed ^33^, the ratio of XY to autosome gene expression starts decreasing in early prophase I, and this decreased ratio is maintained until the late stages of spermatogenesis (Figure 3A). *Ago413^-/-^* males have an increased number of germ cells with upregulated XY genes (Figure 3A) and substantially more upregulated genes on the X and Y compared to the autosomes (Figure 3B). We confirmed the upregulation of XY transcripts in bulk RNA-seq, using sorted pachytene/diplotene spermatocytes and round spermatids (Figure 3C). This pattern of upregulated genes manifests from leptotene-zygotene spermatocytes to post-meiotic spermatids, indicating that AGO proteins are needed for preferential silencing of sex chromosomes during both meiosis and spermiogenesis. To examine MSCI defects in *Ago413*^-/-^ males further, we examined RNA Polymerase II (RNAPII) signal on pachytene spermatocytes, in which RNAPII is usually excluded from the sex body. Notably, *Ago413*^-/-^ males show an increase in RNAPII on the X and Y, compared to both *Ago413*^+/+^ males, (Figure 3D, p <0.01) and *Ago4^-/-^* males ^17^ (35% in triple knockouts vs 18% in single knockout ^17^). Together, these results indicate that AGO3 and AGO4 play roles in MSCI induction and maintenance.

**Figure 3.**
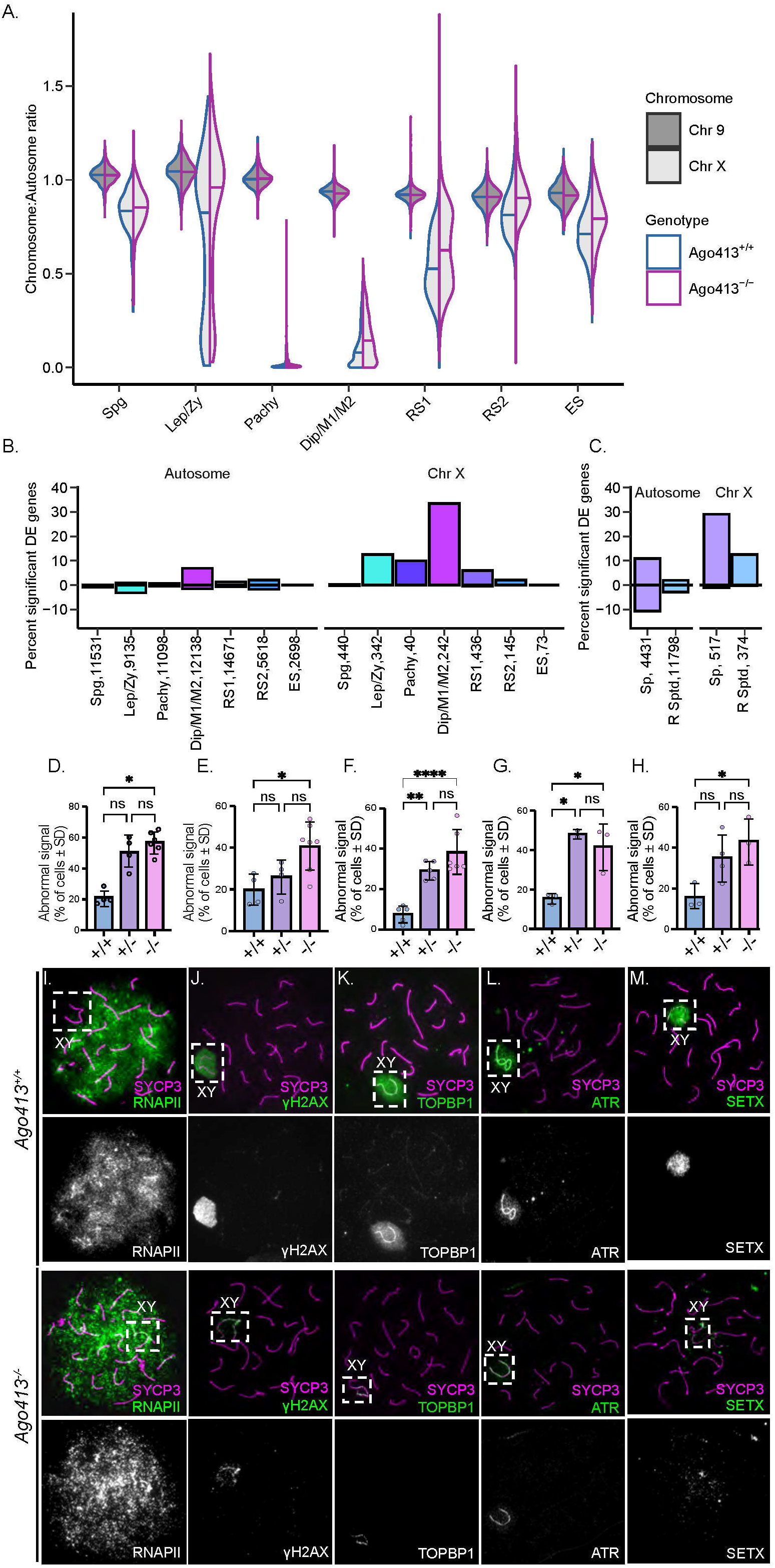
Defective sex chromosome inactivation in *Ago413^-/-^* germ cells. (A) Violin plots displaying the ratio of the average expression of either X chromosome genes or chromosome 9 genes to the mean expression of all autosome genes at different stages of spermatogenesis. Horizontal bar indicates the median for each cell population. (B) Bar plot showing number of genes across cell types called as significantly differentially expressed with adjusted p < 0.05 and absolute value log2FC >0.5. Percentage above 0 indicates upregulated genes and percentage below 0 indicates downregulated genes. Numbers next to each cell population in the X axes indicate the total number of tested genes. (C) As in (B) for bulk RNA-seq. Sp: Spermatocyte, R. Sptd: Round spermatid (D-H) Quantification of aberrant localization for different proteins involved in MSCI, RNA Pol II (D), gH2AX (E), TOPBP1 (F), ATR (G), and SETX (H), in *Ago413^+/+^* and *Ago413^-/-^* pachytene spermatocytes and (I-M) representative patterns for normal (*Ago413^+/+^*) and aberrant (*Ago413^-/-^*) localization in chromosomal spreads. Bars represent mean ± SD, dots represent each replicate. Data were analyzed by ANOVA followed by Tuckey test for multiple comparisons. (*p <0.05, **p <0.01, ****p <0.0001).

To further investigate the role of AGO proteins in MSCI, we analyzed the localization of DDR pathway components involved in MSCI initiation and progression. We found that 40% of spermatocytes in *Ago413*^-/-^ males exhibit altered localization of γH2AX, TOPBP1 and ATR (Figure 3E-G, p <0.05, p <0.0001 and p <0.05 respectively). Similarly, the helicase Senataxin (SETX), a target of TOPBP1 ^33^ that is required for remodeling of the X and Y chromatin role during meiosis ^42,43^, also shows reduced localization to the sex body of *Ago413*^+/-^ and *Ago413^-^*^/-^ spermatocytes compared to *Ago413*^+/+^ cells (48 and 41% respectively, vs 15%, p <0.05, Figure 3H). However, these defects in DDR localization to the sex body are not accompanied by crossover or synapsis alternations in *Ago413*^-/-^mice, as evaluated using MLH1 and SYCP1 staining, respectively (Figure S6 A, B). Analysis of chromosome spreads and single-cell transcriptomics suggests that not all pachytene spermatocytes fail to localize DDR factors to the sex body in *Ago413^-/-^* males. Presumably, the defective cell population is preferentially subjected to apoptosis triggered by the MSCI checkpoint ^38,39^, while cells without DDR defects are more likely to progress to diplotene and ultimately produce spermatozoa.

### Post-meiotic spermatids from Ago413^-/-^ males exhibit defective spermiogenesis and poor spermatozoa function

Given that a fraction of *Ago413^-/-^* prophase I cells evade apoptosis despite MSCI defects, we next evaluated the functionality of the resulting sperm. The morphological and physiological changes that underlie spermiogenic differentiation require gene expression programs that drive acrosome and flagellar formation, nuclear compaction and histone to protamine exchange ^44^. Since spermiogenesis defects are reflected in the ability of sperm to fertilize the oocyte, we evaluated the quality of spermatozoa obtained from the cauda epididymis of *Ago413^-/-^* males. We found that sperm morphology is impacted in *Ago413^-/-^* males (p <0.0001), with increased head and tail malformations (Figure 4A, B, p <0.001 for both comparisons). Additionally, *Ago413*^-/-^ sperm exhibit reduced motility (Figure 3C, p<0.01) and ability to fertilize oocytes *in vitro* (Figure 4D, p<0.01).

**Figure 4.**
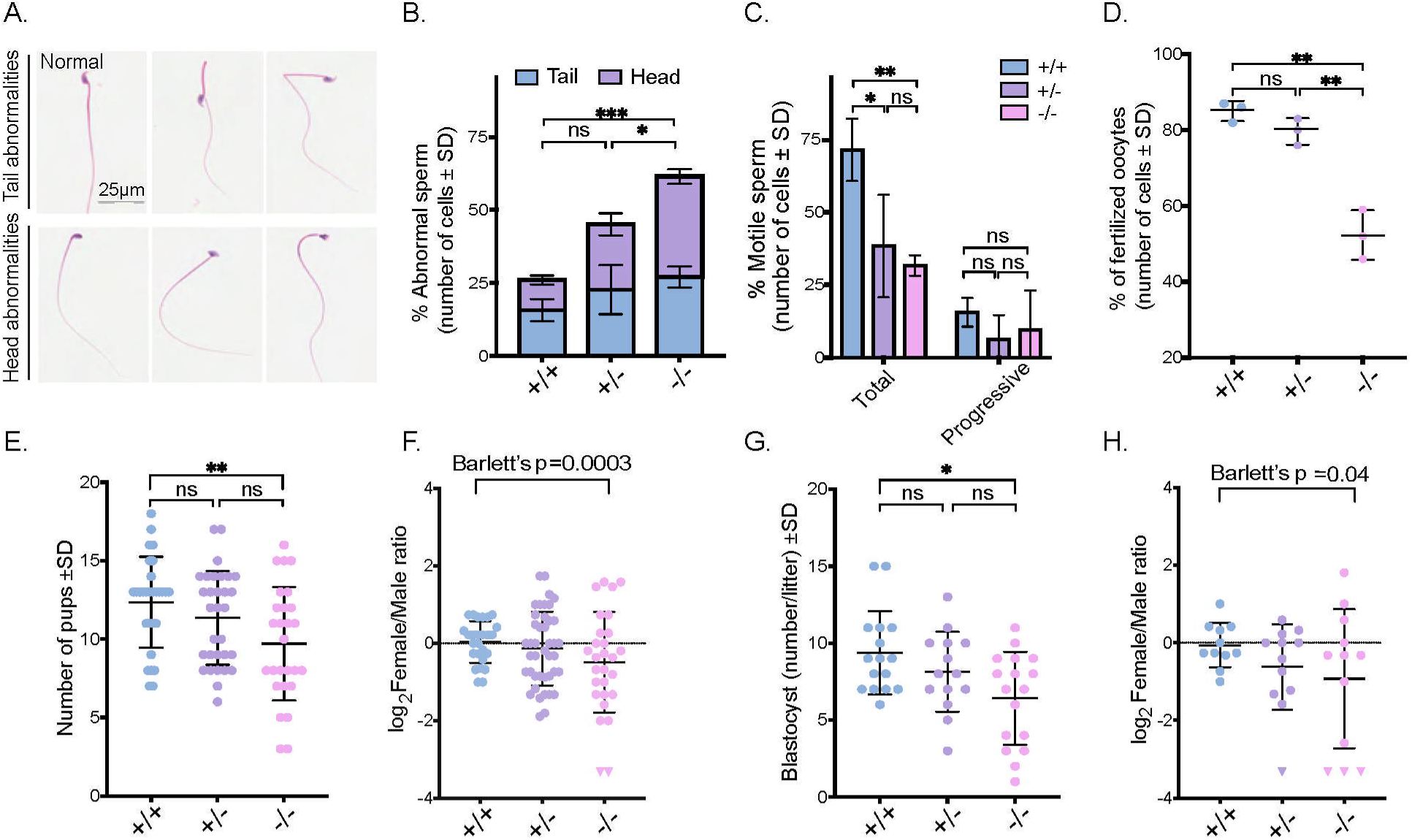
Reproductive phenotype of *Ago413^-/-^* mice. (A) Examples of sperm head and tail morphological abnormalities, normal, bent tail, kinked tail, head tilt, abnormal head shape and lack of acrosome (order as they appear in the figure). (B) Percentage of morphological abnormalities for spermatozoa obtained by swim-up for each genotype. Bars represent each mean value ± SD, n=3. Data were analyzed by Kuskall-Wallis test (*p <0.05, ***p <0.001). (C) Sperm motility parameters obtained by CASA, otherwise as described in panel B (*p <0.05, **p <0.01). (D) Percentage of fertilized oocytes by IVF for each genotype. Dots represent each mean value ± SD, n=3. Data were analyzed by Kuskall-Wallis test (*p <0.05, **p <0.01). (E) Litter size and log2 of female/male pups ratio (F) after natural mating. Dots represent each individual data value and bars represent the mean ± SD. Triangles represent pseudo values for female/male =0. Data were analyzed by ANOVA followed by Tuckey test for multiple comparisons (**p <0.01). Equality of variance of sex ratios data were statistically different according to Barlett’s test (****p <0.001). (G) Number of blastocysts collected at day 3 post-*coitum and* log2 of female/male blastocyst (H) obtained after natural mating. otherwise as described in panels E and F (*p <0.05). Equality of variance of sex ratios data were statistically different according to Barlett’s test (*p <0.05).

We next asked whether the abnormal gametes produced from spermatocytes with defective MSCI and spermatids with reduced post-meiotic sex chromosome silencing in *Ago413*^-/-^ males result in selective X- or Y-bearing spermatid fertilization success. We performed timed mating experiments using wild-type CD1 females, due to the high numbers of oocytes ovulated per estrous cycle by this strain. *Ago413^-/-^* males produce smaller litter sizes (p <0.001) with a larger variation in the sex ratio of pups compared to wild-type mice (p <0.001, Figure 4E, F). To evaluate the developmental stage at which the decrease in litter size and sex ratio variation manifests, we quantified sex ratios at the blastocyst stage. We recovered a lower number of blastocysts from matings between CD1 females and *Ago413^-/-^* males than from matings with wild-type males (Figure 4G, p <0.01), with increased variance in sex ratios (Figure 4H, p <0.05), mirroring the weaning data. These results indicate that embryo loss and altered sex ratios occurs prior to implantation, rather than being the result of embryonic wastage during post-implantation development.

To determine if the variation in X and Y bearing sperm from *Ago413^-/-^* males derives from spermatogenic defects, we checked for differences in the proportions of X and Y bearing sperm in the epididymides. We observe no significant alterations in X and Y sperm in either *Ago413^+/-^* or *Ago413^-/-^* males, as measured by digital droplet PCR, which we also verified by DNA FISH (Figure S7, p >0.05). These data suggest that the disrupted spermatogenic program in *Ago413^-/-^* males affects sperm carrying each sex chromosome equally and that the sex ratio variance in *Ago413^-/-^* progeny is not due to preferential loss of X or Y-bearing sperm. Taken together, *Ago413^-/-^* animals exhibit reductions in sperm motility parameters and fertilization capabilities, accompanied by alterations in the sex ratio of their offspring, the latter arising after sperm production but prior to blastocyst formation.

### Impact of loss of AGO1, 3 and 4 on microRNA-mediated regulation during spermatogenesis

To better understand the extent to which germline phenotypes in *Ago413^-/-^* males are explained by the canonical function of Argonaute proteins with miRNAs, we first investigated which miRNAs are found in the germline and how their expression changes in *Ago413*^-/-^ animals. We performed small RNA-seq using enriched pachytene/diplotene spermatocyte and round spermatid cell fractions and identified annotated mature miRNAs with high or statistically different counts between *Ago413*^-/-^ and *Ago413*^+/+^ males (Figure S8A, B; Table S3, S4). Next, we took these identified miRNAs, as we expect miRNA which have high counts or statistically different counts to show the most changes in miRNA mediated regulation, and then systematically examined our transcriptomic data for evidence that miRNA-mediated regulation by these miRNAs is altered in the *Ago413*^-/-^ germline. Thus, we investigated the expression of stringently predicted ^45^ miRNA target genes for these miRNAs, comparing expression in *Ago413*^-/-^ and wild-type cells. We observed only small changes between such targets and background genes without target sites in both our scRNA-seq (Figure S8 C, D) and the RNA-seq data (Figure S8E). Notably, these patterns of mRNA expression do not strongly correlate with changes in miRNA expression. Taken together, this approach suggested there was only a minimal impact on miRNA-mediated regulation in the absence of AGO3 and AGO4, likely due to the high expression of AGO2.

To determine if the upregulation of spermiogenesis genes seen in *Ago413^-/-^* males is due to a lack of miRNA-mediated repression, we explored whether such genes were plausibly regulated by miRNAs and whether this regulation was altered in *Ago413*^-/-^ males by restricting our expression comparisons to just these gene sets. Diplotene spermatocytes (Figure S9A) and bulk spermatocytes (Figure S9B) of *Ago413*^-/-^ males show increased levels of miRNA target repression for genes related to spermatid development, which is not consistent with loss of miRNA targeting being the main driver of early upregulation of spermiogenesis genes. We similarly checked our observed XY gene upregulation in *Ago413^-/-^* males to determine if this upregulation was caused by a reduction in miRNA mediated regulation and see that miRNA target genes on the XY also do not show expression changes consistent with a loss of miRNA targeting (Figure S9C, D). We confirmed the results of our comparison of expression between all miRNA targets and background genes using an orthogonal miRNA target detection method and see only a small number of miRNAs that are significantly associated with changes in target mRNA levels in spermatogonia through pachytene spermatocytes (Figure S9C-E), and no significant associations after pachynema. Given the lack of large, consistent changes in miRNA targeting signatures or de-repression of spermiogenesis genes or XY-linked genes driven by loss of miRNA regulation, these data suggest that direct canonical, cytoplasmic miRNA regulation alone does not explain the disrupted gene regulation we observe in the germline of *Ago413^-/-^* males, an interpretation consistent with the nuclear localization of AGO3 and AGO4 in the germline.

### Identification of AGO3 and AGO4 interacting proteins in the germline

To gain insights into the mechanisms by which AGO proteins regulate gene expression during meiosis, we identified AGO3 and AGO4 interacting proteins by immunoprecipitation followed by Mass Spectrometry (MS, Figure S10). AGO3 was immunoprecipitated from germ cell lysates of *Ago3*^myc-flag^ mice using anti-MYC conjugated magnetic beads; as a negative control, we repeated this analysis using instead a wild-type lysate (Figure S10 A). Similarly, to identify AGO4 interacting proteins, we utilized an antibody against endogenous AGO4, employing anti-IgG antibodies to generate a background profile (Figure S10 B). We performed three biological replicates for each of the four conditions and calculated an abundance ratio between each AGO IP and respective control (Table S5, S6). We selected candidate interactors with consistent enrichment in at least two replicates. Notably, proteins we identified as AGO3 and AGO4 interactors are predicted to be localized to the nucleus (76% and 72%, respectively; Figure S10C, D). Gene ontology analysis of these candidate interacting proteins indicates enrichment in Biological Process terms associated with small RNA pathways and mRNA metabolism, for both AGO3 and AGO4 (Fig S10E, F). Interestingly, the overlap between AGO3 and AGO4 candidate interactors is low, with only the RNA binding protein HNRNPD ^46^ and PPP5C, a phosphatase that regulates the ATM/ATR signaling during the DDR ^47^, shared between AGO3 and AGO4.

AGO3 and AGO4 also have largely non-overlapping candidate nuclear interactors. AGO3-specific interactors include proteins associated with chromatin, including CTCF, the cohesin stabilizer MAZ ^48^, the H4 interacting protein MEPCE ^49^, and BRG1, the ATPase domain of the chromatin remodeling BAF complex ^50^. In contrast, candidate nuclear interacting proteins for AGO4 include the catalytic core component of RNA polymerase II, POLR2B, as well as YBX2, an RNA-binding protein expressed in pachytene spermatocytes and round spermatids, which binds to promoters in addition to regulating translation ^51^. Overall, binding partners of AGO3 and AGO4 confirm their functionality within the nucleus of the germ cell and suggest that their role involves interaction with the transcription and chromatin remodeling machinery.

### AGO mediated gene regulation in the germline

Our data indicate a non-canonical role for nuclear AGO3 and AGO4 in the germline. Therefore, we sought to identify transcription factors which may be involved in the changes in gene expression observed in the *Ago314*^-/-^ germline. We first identified candidate regulatory regions proximal to significantly de-repressed genes in *Ago413^-/-^* cells compared to wild-type cells in spermatocyte clusters identified in our scRNA-seq data. These regions were previously annotated as active enhancers using a combination of nascent RNA-sequencing and genome-wide mapping of open chromatin applied to leptotene/zygotene, pachytene and diplotene spermatocytes ^52^. We then performed transcription factor motif enrichment ^53^ on the enhancer regions proximal to genes whose expression changes in *Ago314*^-/-^ cells ^56^. Evaluation of regulatory regions near all upregulated genes in the diplotene to M2 cluster of the scRNA-seq data set reveals that 5 of the top 10 most enriched motifs correspond to the AP-1 TF family (Figure 5A, Table S7), enrichment that we also observed when we examined only XY genes (Figure 5B, Table S8). Moreover, we confirmed AP-1 TF motif enrichment when we analyzed regions associated with upregulated genes in the bulk RNA-seq of spermatocytes (Figure 5C, Table S9). We evaluated expression of AP-1 factors in our scRNA-seq (Figure S11A) and used immunofluorescence on testicular sections to examine the staining patterns of the AP-1 factors JunD, ATF2 and ATF7 in the germline (Figure S11B). Strikingly, ATF2, one of the enriched AP-1 motifs in the XY upregulated genes, shows localization to the sex body in wild-type spermatocytes (Figure 5D).

**Figure 5.**
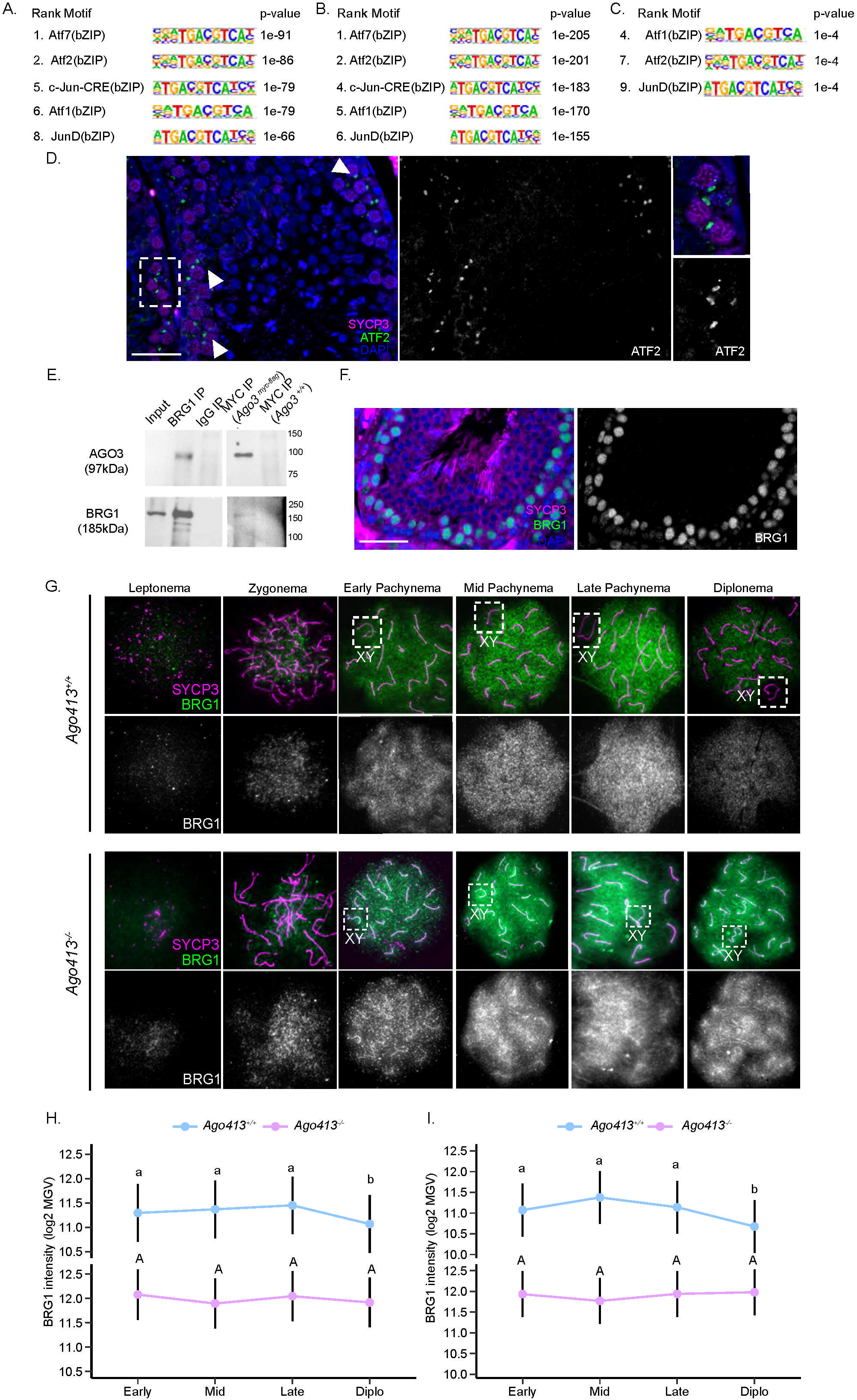
Identification of AGO3 and AGO4 mediated gene regulation through AP-1 factors. (A- C) AP-1 factors found by HOMER motif enrichment analysis results of regulatory regions within 2kb of significantly upregulated genes in *Ago413^-/-^* germ cells compared to wild-type cells in (A) scRNA-seq analysis of the Dip/MI/MII cluster, (B) specific XY-linked genes upregulated in diplotene scRNA-seq analysis, and (C) bulk RNA-seq of pachytene/diplotene enriched fractions. (D) Immunostaining of *Ago413^+/+^* testicular sections with ATF2 and SYCP3 antibodies, showing localization of ATF2 to the sex body of pachytene spermatocytes. (E) Western blot showing co-immunoprecipitation of AGO3 and BRG1 in *Ago3^myc-flag^* germ cells. (F) Immunostaining of *Ago413^+/+^* testicular sections with BRG1 and SYCP3 antibodies, showing localization of BRG1 to the pachytene spermatocyte population. (G) Prophase I spreads of *Ago413^+/+^* and *Ago413^-/-^* spermatocytes stained with SYCP3 and BRG1 showing localization patterns for BRG1. (H-I) Log2 of Mean gray value of BRG1 fluorescence intensity per cell quantified using a macro script for Fiji as previously reported ^52^ for each substage of prophase I in whole cell (H) and only the sex chromosomes (I) in meiotic spreads from *Ago413^+/+^* and *Ago413^-/-^* testes. Points represent estimated marginal mean for all samples of each genotype; different colors represent genotype and vertical line represents 95% confidence interval. Data were analyzed by fitting mixed linear model with random effect for sample and calculation of estimated marginal means followed by pairwise Tukey test (n=4, *p <0.05, **p <0.01, ***p <0.001). Different letters represent a statistical difference between groups.

AP-1 TFs have been widely characterized for their regulation capacities during cell proliferation ^54^ and are known to interact with the BRG/BRM-associated factor (BAF) complex, a chromatin remodeling complex, to promote accessibility of binding sites for other TFs ^55^. This chromatin remodeling is mediated by the ATPase domain of the BAF complex, BRG1, ^56–60^, which we identified as a candidate AGO3 interactor (Figure S10C). We validated this interaction by reciprocal co-IP assays of BRG1 and AGO3 (Figure 5E). To examine BRG1 protein through spermatogenesis, we performed immunofluorescence staining in testis tubules of wild-type mice (Figure 5F) and observed that BRG1 is localized to the spermatogonia-prophase I layer of the seminiferous tubules and is not present in spermatids onwards, suggesting a role during the meiotic stage of spermatogenesis, consistent with previous studies ^61^.

To examine the link between AGO3 and BRG1 further, we examined BRG1 localization in prophase I spreads from *Ago314*^-/-^ males and wild-type littermates and observed that BRG1 signal is low in leptonema and zygonema and becomes brighter during pachynema (Figure 5G). In wild type spermatocytes, BRG1 covers the whole XY chromatin domain by early pachynema, before becoming progressively excluded by diplonema (Figure 5G). Total and XY BRG1-associated fluorescence intensity show a reduction in the transition from late pachynema to diplonema in wild-type spermatocytes, while they remain constant along all substages in *Ago314*^-/-^ spermatocytes, leading to higher levels of BRG1 in diplotene spermatocytes in the absence of AGO3 and AGO4 due to lack of removal of BRG1. Thus, our data point to a role for AGOs in removal of BRG1 from the chromatin after the sex body has been formed.

## DISCUSSION

The Argonaute family of proteins is essential for miRNA-mediated post-transcriptional gene regulation ^21^. The extent to which mammalian AGOs function in the nucleus, or independently of miRNAs, to impact transcription and transcription-related events is unclear, despite extensive studies in *S. pombe* and *C. elegans* indicating roles for AGOs in co-transcriptional repression and recruitment of repressive histone marks ^22–26^. In mammals, AGO3 and AGO4 localize to the heterochromatin-rich sex body of mouse spermatocytes ^17^, suggesting a role for these proteins in transcriptional silencing of the sex chromosomes. Here, we have used immunofluorescence and single-cell transcriptomics to extend upon these findings, revealing complex and distinct localization patterns for AGO2, AGO3 and AGO4 during spermatogenesis. *Ago1* is predominantly expressed in spermatogonia and not later stages, while *Ago2*, *Ago3* and *Ago4* are expressed in prophase I spermatocytes. At the protein level, AGO2 is predominantly cytoplasmic, while AGO3 and AGO4 have a distinct nuclear localization, supporting functional partitioning of AGO proteins in the male germline. AGO3 localizes to the synaptonemal complex of all chromosomes in early pachynema and becomes restricted to the sex body by late pachynema. By contrast, AGO4 localizes to the sex body from early pachynema and persists until at least diplonema. Additionally, we observe AGO4 localization to pericentromeric heterochromatin of all chromosomes in pachytene and diplotene spermatocytes. Interestingly, DICER, which is required to generate small RNAs that partner with AGOs, localizes to these heterochromatic regions in mouse spermatocytes, where it binds to major satellite transcripts ^62^. Our data provide further evidence for a functional association between AGO-dependent small RNAs and heterochromatin formation, maintenance and/or silencing.

The reproductive defects of *Ago413^-/-^* mice show that AGO3 and AGO4 regulate proper timing of gene expression during spermatogenesis and participate in MSCI. Four lines of evidence suggest that these functions of AGO3 and AGO4 are driven by events in the nucleus, rather than a more conventional role in mediating cytoplasmic regulation of miRNA targets. First, identification by western blot in nuclear fractions and immunofluorescence of testicular germ cells confirms the enrichment of these Argonautes in the nucleus of prophase I cells. Second, de-repressed spermiogenesis genes and sex-linked genes are not enriched in miRNA targets of differentially expressed miRNAs in *Ago413^-/-^* males. Third, candidate protein binding partners of both AGO3 and AGO4 are enriched in the nucleus where they play roles in transcription and chromatin remodeling. Lastly, we confirmed an interaction between AGO3 and BRG1, the ATPase subunit of the BAF complex, and observed an accumulation of BRG1 in the nucleus of the spermatocytes when AGO3 is absent. We propose that the role of AGO3 in MSCI involves interaction with the chromatin remodeling machinery though BRG1. AGO4 likely has a synergistic role that is at least partially independent from AGO3 given its different timing of accumulation in the sex body, perhaps related to interactions with POLR2B and/or other transcriptional machinery, which will require further study.

The BRG1, as well as other BAF complex subunits are required for mouse spermatogenesis and MSCI ^61,63–66^. BRG1 particularly, mediates recruitment of the polycomb protein SCML2 and the deubiquitylase USP7 to the sex chromosomes during MSCI ^67,68^. Furthermore, an interaction between AGO proteins and the catalytic subunits of BAF has been described in mammalian cells, including mouse spermatocytes ^30,69^. Our data suggest that BRG1 and AGO3 interact in early pachynema, when AGO proteins localize to the sex body and autosomal SCs and BRG1 is present throughout the nucleus. This occupancy of BRG1 prior to and at the beginning of the transcriptional shutdown in spermatocytes is reminiscent of the increase in chromatin occupancy of BRG1 on the inactivated X chromosome at the beginning of X chromosome inactivation (XCI) in female mammals. At the start of XCI, BRG1 is required to create nucleosome-depleted regions at promoters and is later removed from the chromatin by the lncRNA Xist ^70,71^. We hypothesize that BRG1 localization to the XY at the beginning of pachynema and its subsequent removal by diplonema, when the sex body heterochromatin has been fully established, corresponds to similar chromatin dynamics than XCI, and may also be reliant on RNA interactions. Thus, we propose that persistence of BRG1 in diplotene cells might be associated with the changes in gene expression observed in *Ago314*^-/-^ spermatocytes. In this model, the localization of ATF2 to the sex body indicates that failure to repress XY genes is likely due to a BRG1-ATF2 mediated increase in accessibility of the sex chromosomes, leading to alterations in gene expression which dysregulate spermiogenesis progression and cause phenotypic changes in fertility.

Regardless of the mechanism involved, disruption of gene expression timing during prophase I due to lack of AGO3 and AGO4 directly impacts the germ cell developmental program. A prominent feature of this dysregulation is the switch between the delayed expression of genes required for meiotic progression in pre-pachytene germ cells and premature expression of genes needed in diplonema and later stages. The transition between pachynema and diplonema includes the progression of cells beyond the pachytene checkpoint that coordinates homologous recombination and cell cycle progression ^39^. In male mammals, failure to complete MSCI also leads to pachytene arrest and elimination of spermatocytes by apoptosis ^13,40^. *Ago413*^-/-^ mice exhibit defective MSCI, but only a fraction of pachytene cells that overexpress XY genes die at the pachytene checkpoint, while other spermatocytes are able to complete meiosis and move into spermiogenesis to generate a higher proportion of defective spermatozoa. We hypothesize that the low quality of *Ago413*^-/-^ spermatozoa is due to the early engagement of diplotene spermatocytes in the spermiogenesis program through upregulation of post meiotic genes and defective silencing of sex chromosomes during spermiogenesis, a process known as post meiotic sex chromosome repression ^72–74^. Altogether, this gene dysregulation during spermiogenesis contributes to generation of spermatozoa with a higher frequency of morphological defects, poor motility, and reduced ability to interact with the oocyte. These meiotic and post meiotic phenotypes make the *Ago413*^-/-^ mouse model distinct from other MSCI defective mouse knockouts for genes that function upstream of the ATR-mediated DDR pathway that initiates MSCI, which feature meiotic arrest at the pachytene-diplotene transition ^13,33,41,75,76^. Here we show that AGO3 and AGO4 coordinate gene expression to promote prophase I progression, while the gene dysregulation caused by the lack of these two Argonautes in the post-meiotic phase of spermiogenesis derives specifically from their role in meiotic prophase I. The persistence of sex chromosome de-repression that starts in pachynema and continues until the spermatid stage suggests that AGO3 and AGO4 contribute to the epigenetic silencing that is established during meiosis and which persists until spermiogenesis. This dysregulation makes the *Ago413^-/-^* mouse a unique model to better understand the interrelatedness between prophase I gene regulation and spermatid differentiation. Taken together, these studies demonstrate that AGO3 and AGO4 are key regulators of spermatogenesis, orchestrating expression of meiosis-related genes during prophase I while maintaining silencing of spermiogenesis genes through novel mechanisms in the germline nucleus.

## ACKNOWLEDGEMENTS

This work was supported by funding from the Eunice Kennedy Shriver National Institute of Child Health and Development to PEC, AG and JG (P50HD076210 and P50HD104454), and to PEC (R01HD041012). Additional funding was provided through the Lalor Foundation Postdoctoral Fellowship program and the Cornell Center for Vertebrate Genomics to MC, and the Bill and Melinda Gates Foundation to PEC (INV-035106). We thank the Genomics (RRID:SCR_021727), the Transcriptional Regulation and Expression (TREx, RRID:SCR_022532), the Genomics Innovation Hub (RRID:SCR_022547), the Flow Cytometry (RRID:SCR_021740) and Proteomics and Metabolomics Facilities (RRID:SCR_021743) of the Biotechnology Resource Center of Cornell Institute of Biotechnology for their help with sequencing, cell sorting and proteomics experiments. We want to thank Dr. Joana VIdigal from the Laboratory of Biochemistry and Molecular Biology, National Cancer Institute (NIH) for kndly providing the *Ago2* ha mouse model and Rob Munroe and Chris Abratte from the Stem Cell and Transgenic Core Facility at Cornell for generating the *Ago3* myc flag mouse model. We are grateful to Cornell Center for Animal Resources and Education (CARE) for assistance with mouse husbandry and Eileen Shu and Elizabeth Fogarty for overall lab management as well as the entire Cohen and Grimson labs for feedback and advice during the course of this research.

## Author contributions

Conceptualization: MC, AD, PEC, AG, JKG

Methodology: MC, AD, PEC, AG, JKG

Software: AD, FA, JKG

Validation: MC, AD, PEC, AG

Formal analysis: MC, AD, PEC, AG, JKG, FA

Investigation: MC, AD, ATM, EP, MA, LS, ST

Resources: AG, PEC, JKG

Data Curation: MC, AD, JKG, FA

Writing – Original Draft: MC, AD, AG, PEC

Writing – Review & Editing: MC, AD, AG, PEC, JKG

Visualization: MC, AD, AG, PEC

Supervision: AG, PEC, JKG

Project Administration: AG, PEC

Funding Acquisition: PEC, AG, JG

## SUPPLEMENTARY FIGURES

**Figure S1.**
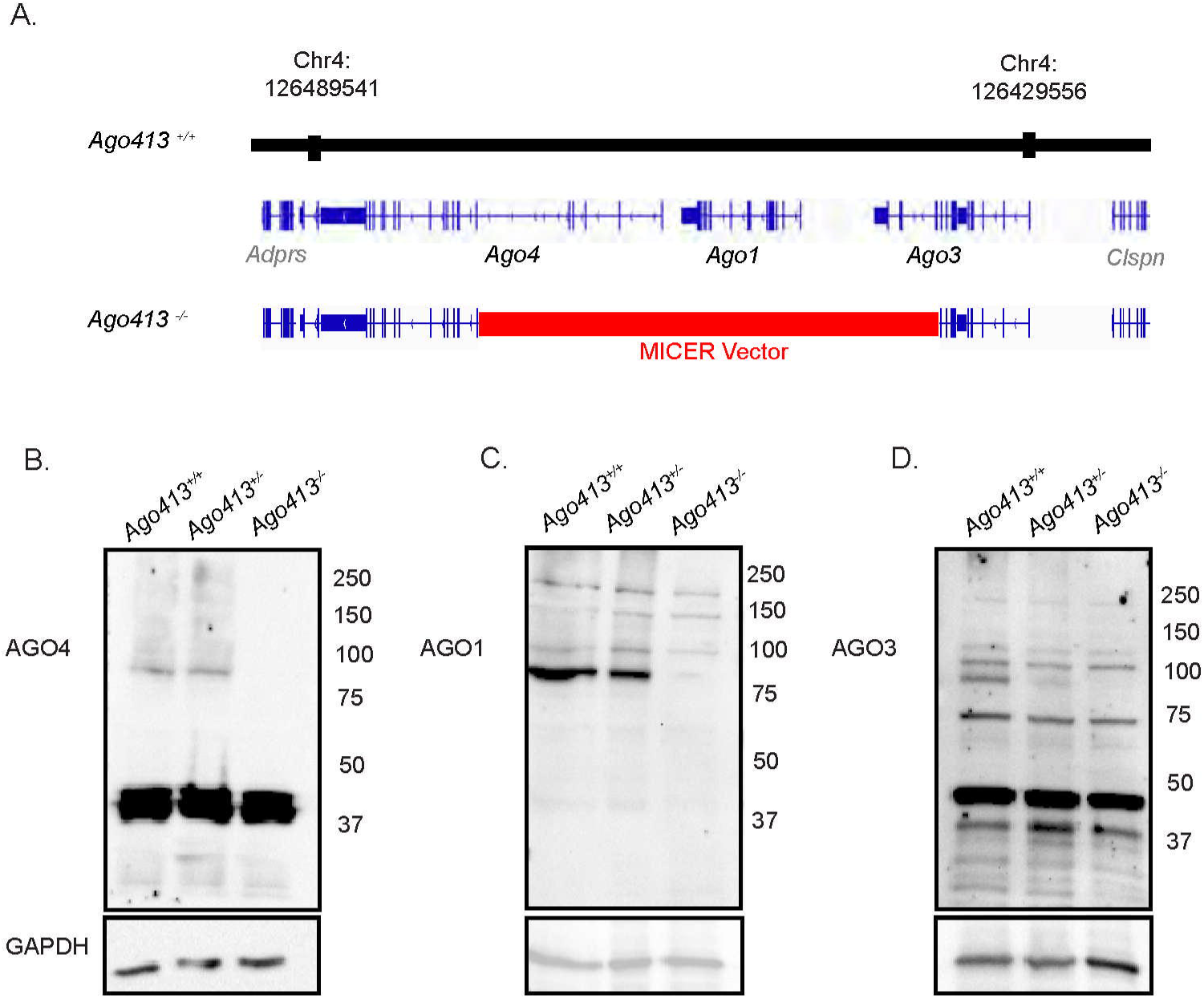
AGO localization patterns during spermatogenesis. (A) *Ago413* knockout mouse model. Western Blots using different anti-AGO antibodies: AGO4 (B), AGO1 (C) and AGO3 (D) on testis lysates from wild-type, heterozygous and null *Ago413* mice. 60 μg of protein was loaded per lane, GADPH was used as a loading control. Expected AGO size (97 kDa) numbers at the right indicate weight marker (kDa).

**Figure S2.**
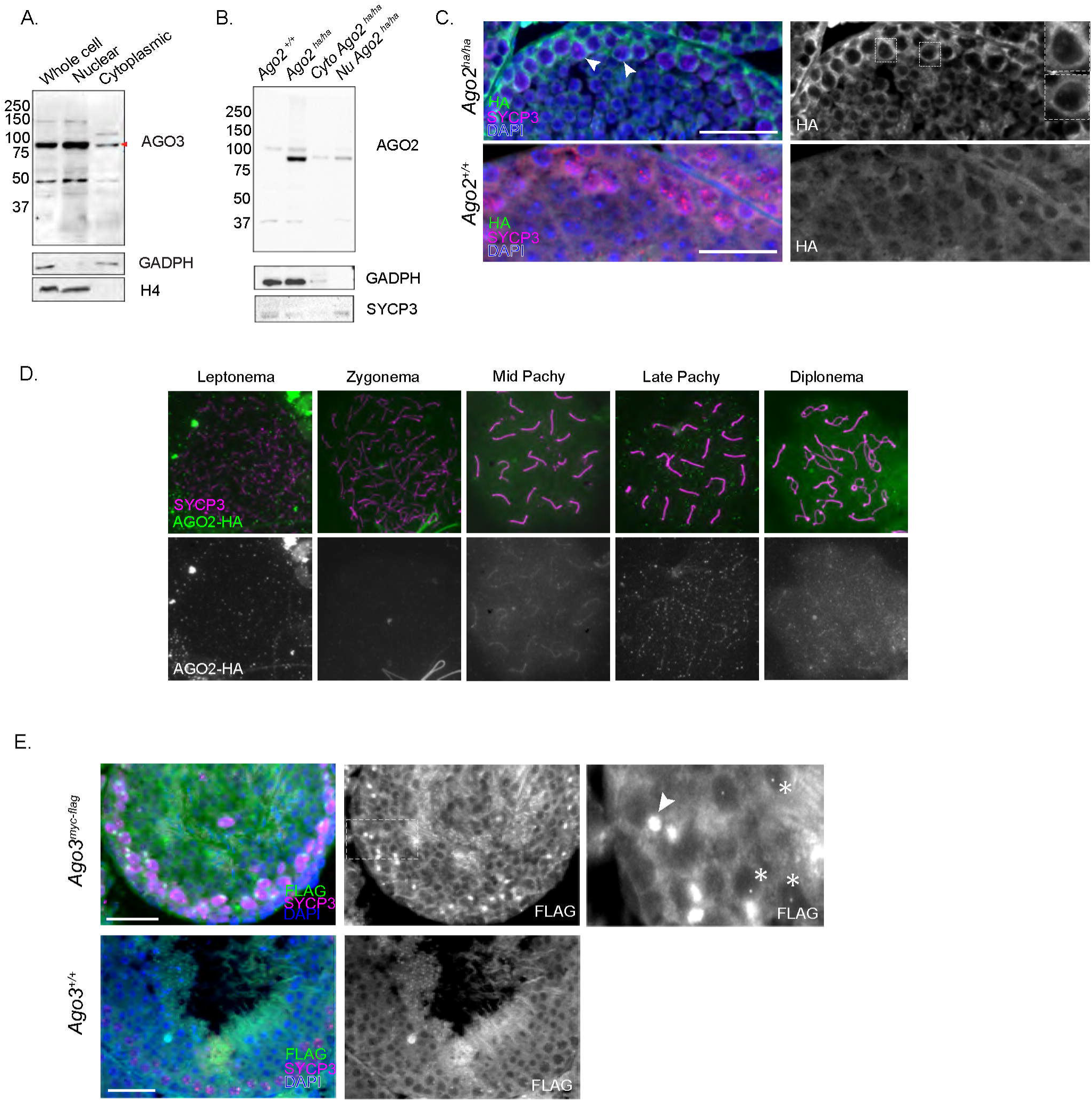
(A) Western Blots using an anti-MYC antibody on germ cell protein lysates from whole cell, nuclear and cytoplasmic fractions of *Ago3 myc flag* homozygous mice. 60 μg of protein was loaded per lane, GADPH and Histone 4 were used as a loading control of cytoplasmic and nuclear fractions respectively. (B) Western Blot with anti-HA antibody in whole cell, nuclear and cytoplasmic protein lysates from *Ago2^ha/ha^* and *Ago2^+/+^* germ cells showing presence of AGO2-HA in both subcellular compartments, GADPH and SYCP3 were used as loading controls of cytoplasmic and nuclear fractions. 60 μg of protein was loaded per lane, expected AGO size (97 kDa) indicated with an arrow. (C) Testicular sections of homozygous *Ago2^ha/ha^* and *Ago2^+/+^* mice immunostained with SYCP3 and HA antibodies and DAPI. Arrows point at presence of AGO2 in the spermatocyte cytoplasm. Enlarged images show lighter staining in the nucleus compared to the cytoplasm. (D) Prophase I spreads of *Ago2^ha/ha^* mice immunostained with SYCP3 and anti-HA antibody. (E) Testicular sections of homozygous *Ago3^myc-flag^* and *Ago3^+/+^* mice immunostained with SYCP3 and FLAG antibodies and DAPI. Arrows in the enlarged images point at presence of AGO3 in sex bodies, while asterisks show localization of AGO3 to round spermatids nucleus. Bars indicate 40 μm.

**Figure S3.**
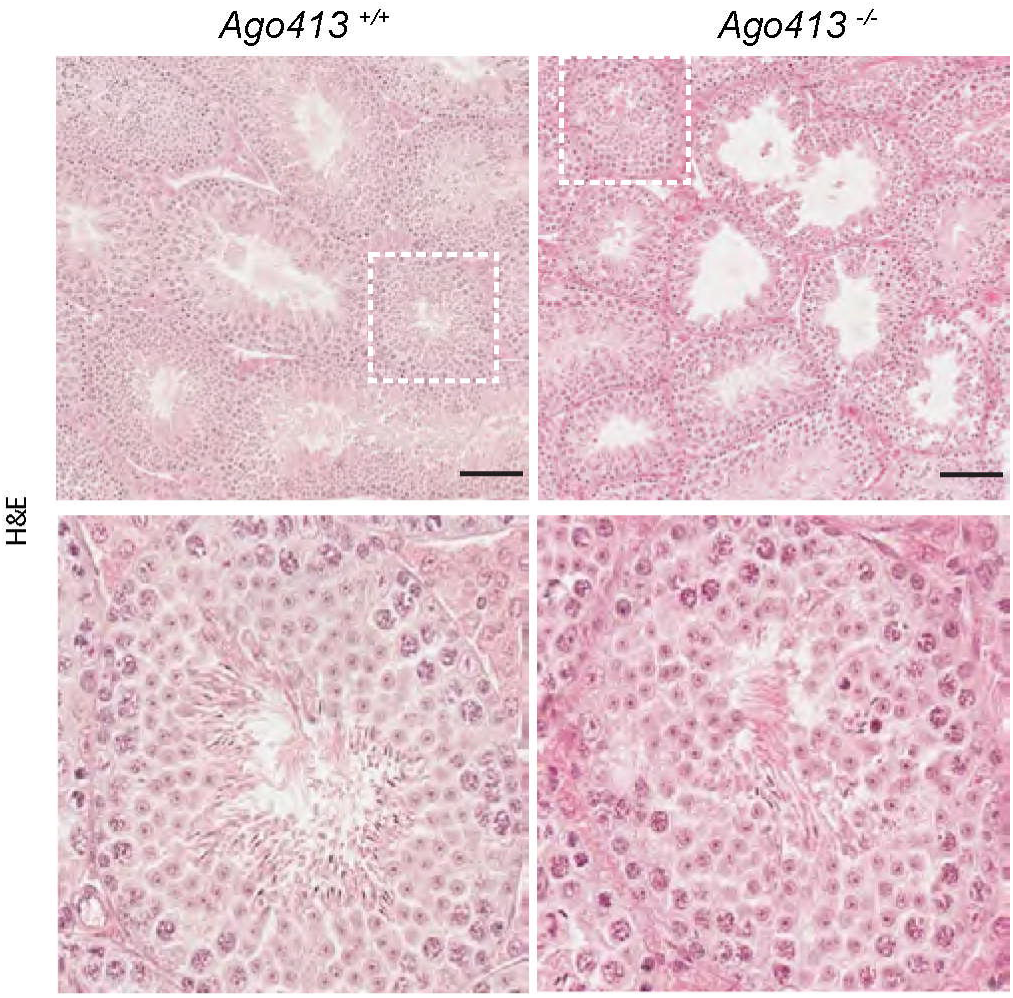
Hematoxylin-Eosin staining of *Ago413^+/+^* and *Ago413^-/-^* testicular sections showing testicular architecture, magnified in panels on the bottom. Bars indicate 40 μm.

**Figure S4.**
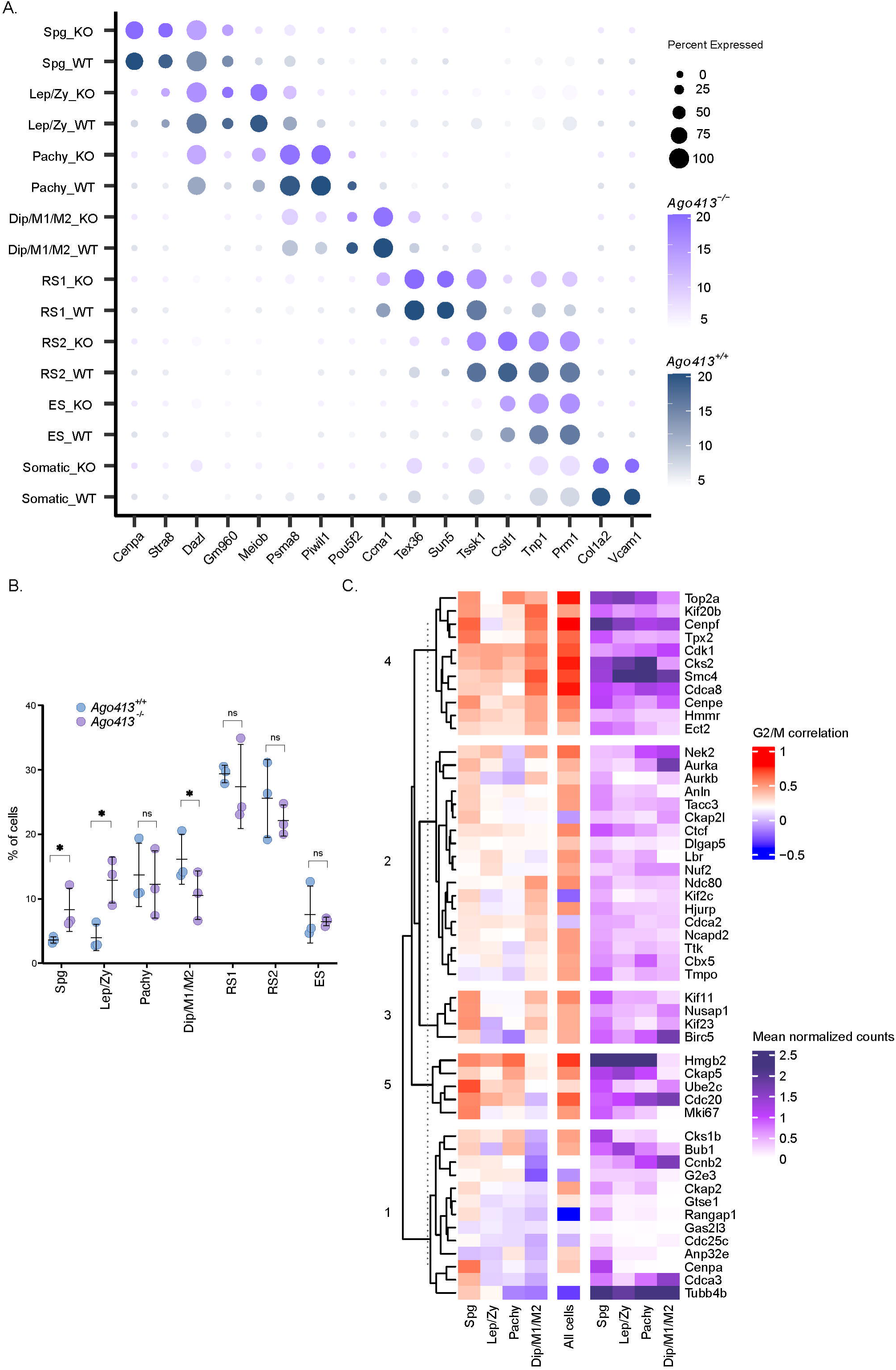
Single cell RNA-seq Supplementary Data. (A) Expression levels of markers of germ and somatic cell types separated by cell type and genotype. (B) Percentage of cells by genotype identified as each cell type by marker gene expression in scRNA-seq. Each dot represents an individual sample. Bars represent the mean ± SD, n=3. Differential proportion p-values tested using sccomp package sum-constrained independent Beta-binomial distribution testing (* p<0.05). (C) Heatmap of Spearman correlation and expression levels of individual G2M marker genes. Correlation was calculated between G2/M score and normalized gene expression for all cells within clusters and for all cells in dataset. Expression levels are mean normalized counts per cell for each cluster.

**Figure S5.**
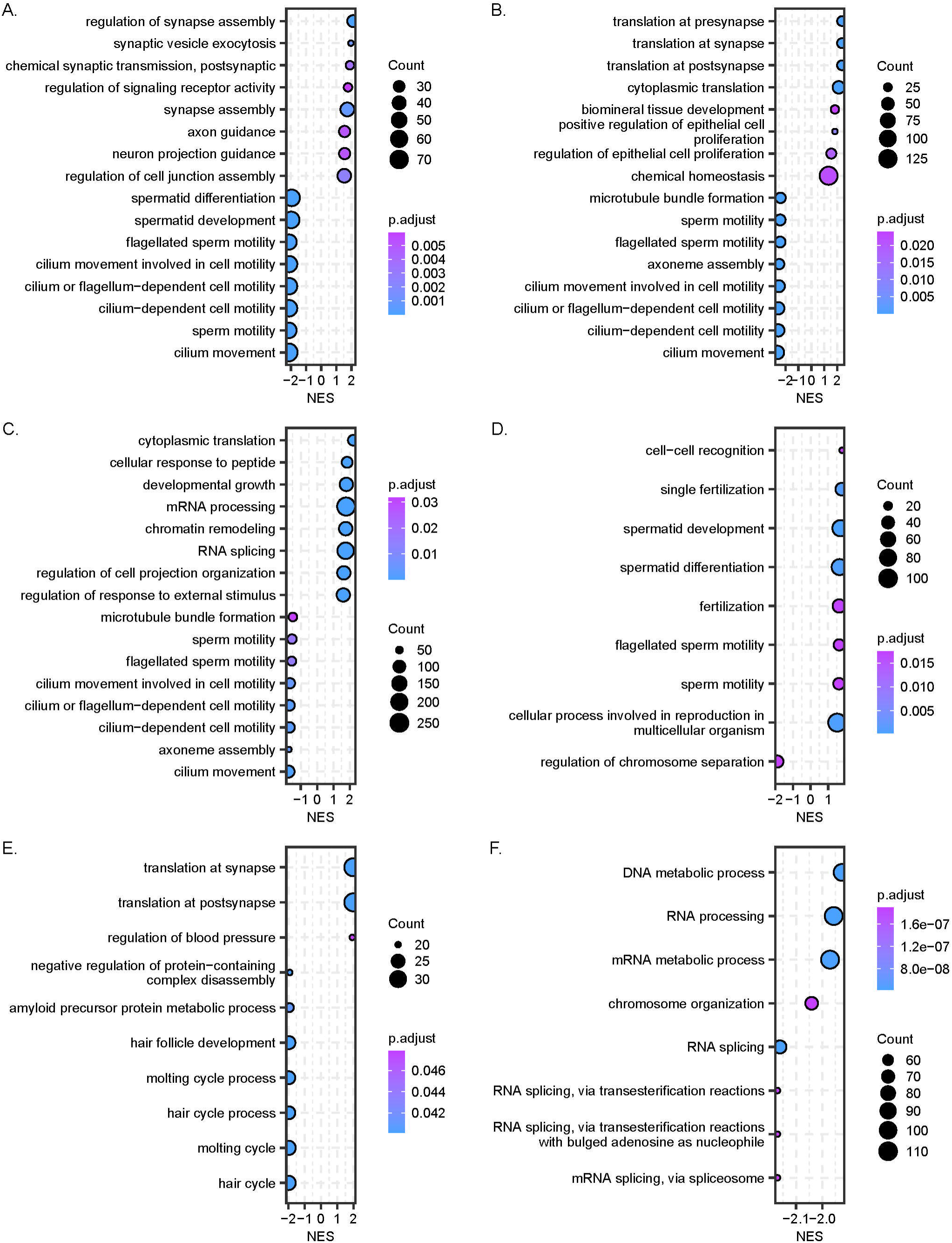
Gene Ontology gene set enrichment analysis. Dot plot showing normalized enrichment score (NES) of biological process gene ontology term gene set enrichment analysis by single cell cluster: (A) Spg, (B) Lep/Zy, (C) Pachy, (D) Diplo/M1/M2, (E) RS1 and (F) RS2. Log2FC values were produced by pseudo bulk differential expression analysis between genotypes and p-values are FDR adjusted.

**Figure S6.**
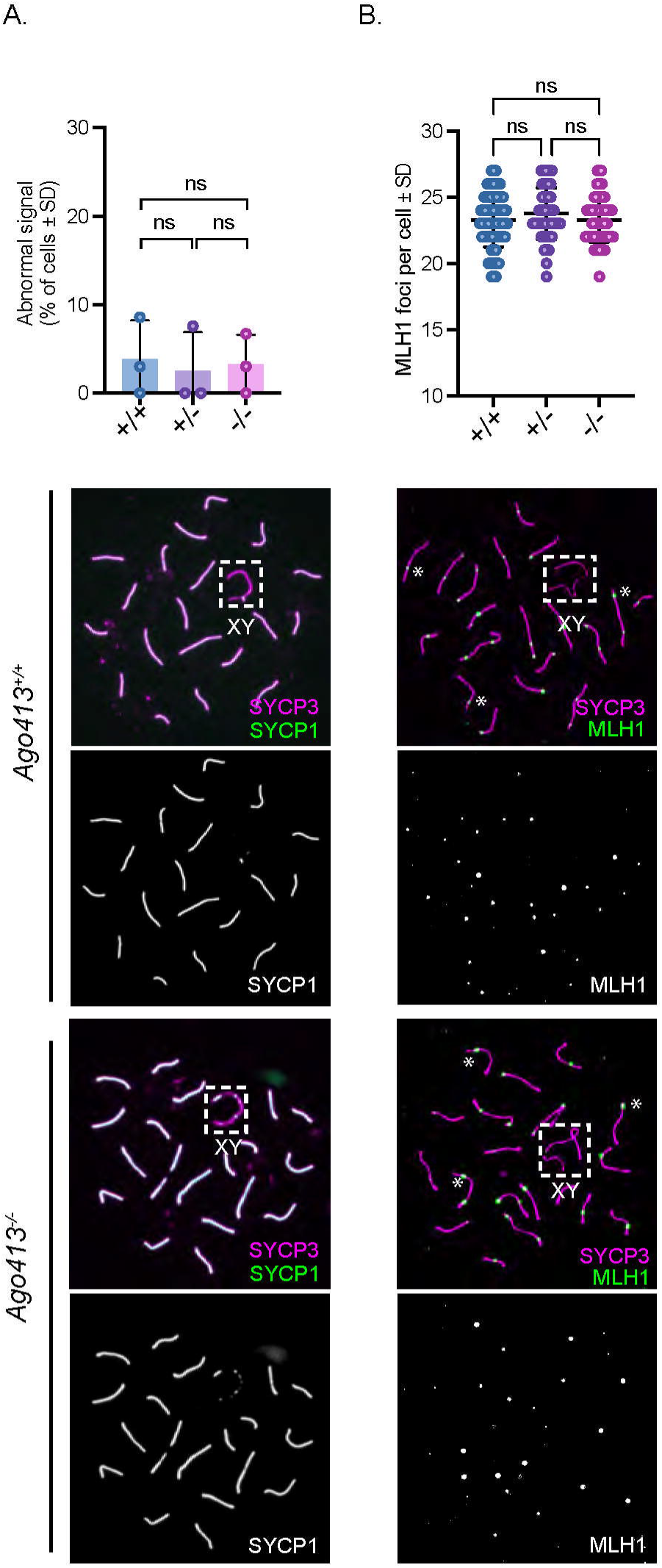
Synapsis and MLH1 counts analysis of the *Ago413* mouseline. Staining patterns and quantification of SYCP1 (A) and MLH1 (B) in pachytene spermatocytes. Bars represent mean ± SD, dots represent each replicate. Data were analyzed by Kruskall-Wallis followed by Dunn’s test for multiple comparisons.

**Figure S7.**
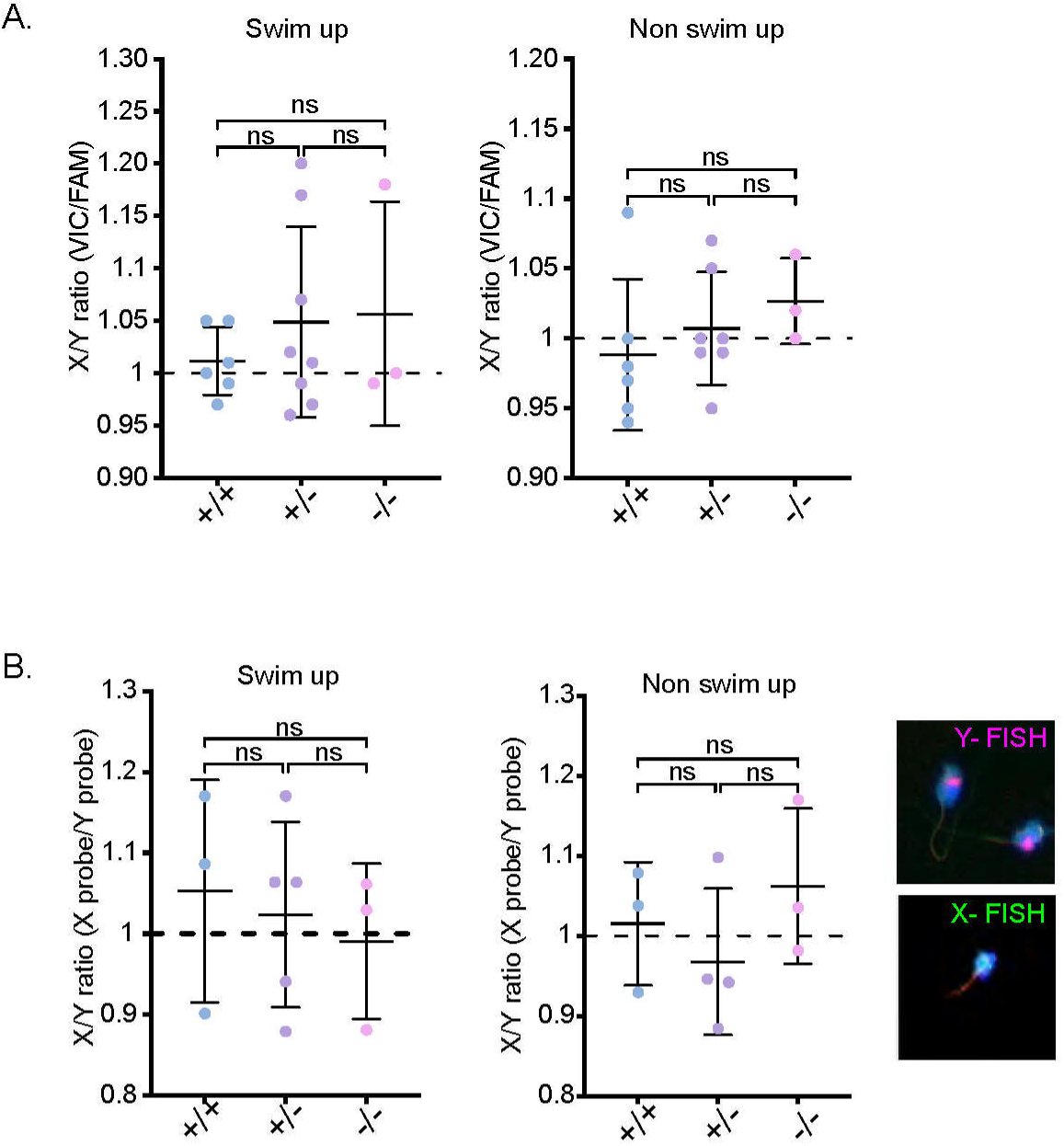
Ratio of X/Y bearing spermatozoa in *Ago413^-/-^* mouse line. (A) Digital droplet PCR of sperm DNA from different *Ago413* genotypes performed in swim up (motile) and non-swim up (immotile) fractions of epididymal spermatozoa. To target the Y and X chromosome, TaqMan Gene Expression Assay probes against the genes *Sry* with a FAM label and *Rbmx* with a VIC label were used respectively. Dots represent the ratio of VIC to FAM signal for each sample ± SD. Data were analyzed by ANOVA followed by Tuckey test for multiple comparisons. (B) Sexing of sperm by targeting the X and Y chromosome for Fluorescent in situ hybridization (FISH). Dots represent the ratio of the number of X-fluorescent sperm over Y-fluorescent sperm for each replicate ± SD. Data were analyzed by ANOVA followed by Tuckey test for multiple comparisons. Example image of fluorescent sperm for X and Y probes stained with DNA dye DAPI.

**Figure S8.**
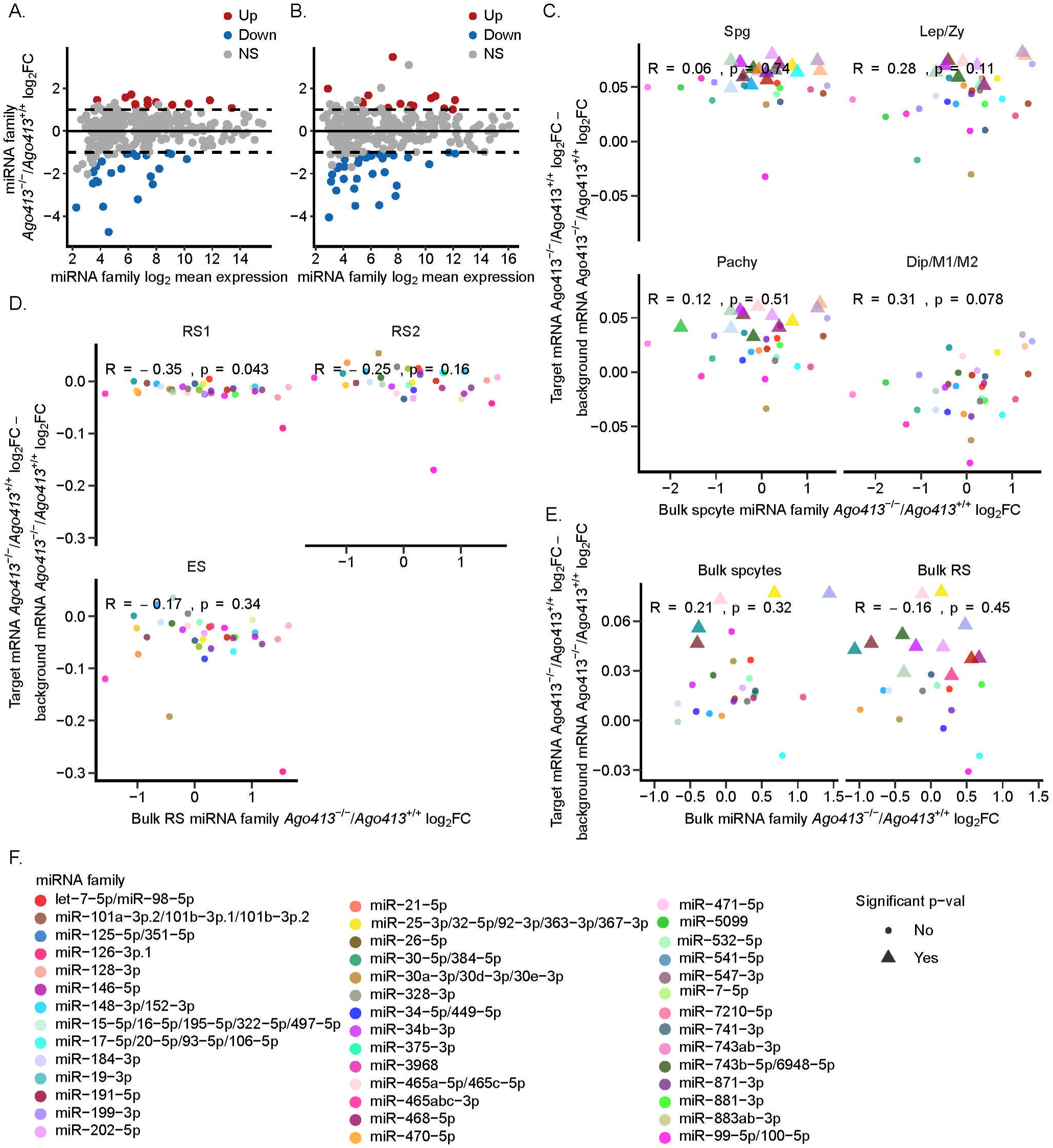
Posttranscriptional gene regulation in germ cells from *Ago413^+/+^* and *Ago413^-/-^* males. (A) MA plot for mature miRNA families identified in smRNA-seq for enriched spermatocytes. Significantly differentially expressed families have an absolute value log2 fold-change greater than or equal to 1 and an adjusted p-value less than 0.05. (B) As in (B) for round spermatids. (C) Scatter plot showing difference in mean log2 fold-change for miRNA targets (Targetscan cumulative weighted context score < -0.2) minus background genes on y-axis and log2 fold-change of miRNA family in bulk spermatocytes on x-axis. Color and shape legend in (F). RNA log2 fold-changes from pseudo bulk analysis of spermatogonia, leptotene/zygotene, pachytene, and diplotene and dividing cell types. Shape and size indicate if multiple test corrected p-value for Wilcoxon rank-sum test between miRNA target log2 fold-changes and background log2 fold-changes are less than 0.05. Spermatogonia through pachytene show small but significant upregulation of gene expression. Diplotene and dividing show no trend towards upregulation. (D) As in (C) for round spermatids 1 and 2 and elongating spermatids, with log2 fold-change of miRNA families coming from bulk round spermatid smRNA-seq. Round spermatid and elongating spermatid clusters don’t show pattern of upregulation of gene expression. (E) As in (C) for bulk RNA-seq analysis. miRNA log2 fold-change comes from corresponding bulk smRNA-seq dataset. Both spermatocytes and round spermatids show small but significant upregulation from some miRNAs.

**Figure 9.**
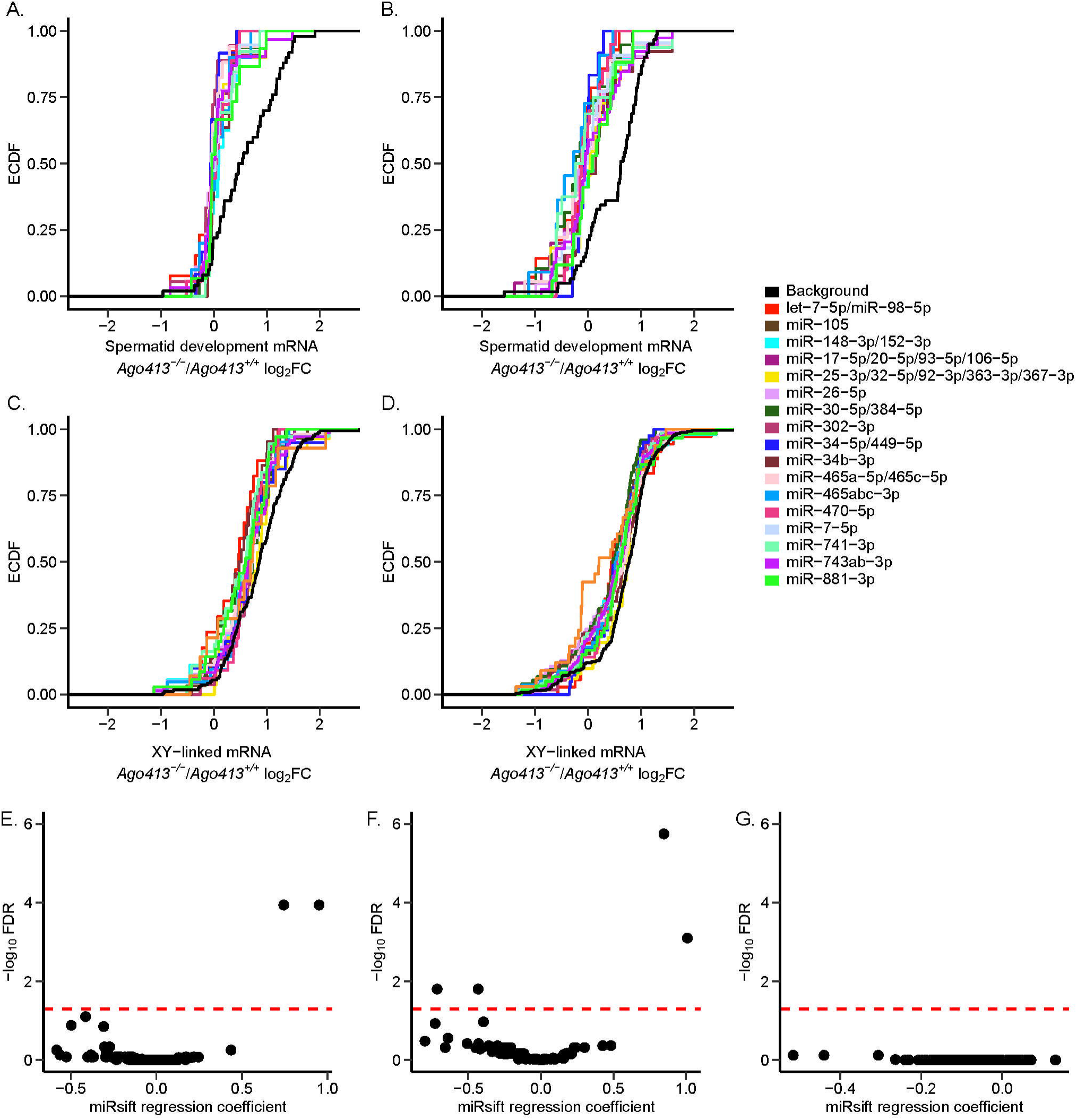
Posttranscriptional regulation in Ago413 spermatogenesis. (A-B) Cumulative distribution plot of KO/WT log2 fold-change from diplotene pseudo bulk analysis (A) and bulk spermatocyte RNA-seq (B), with miRNA targets and background genes subset to only genes in GO term spermatid development. (C-D) As in (A-B) for XY-linked genes for diplotene pseudo bulk analysis (C) and bulk spermatocyte RNA-seq (D). (E) Scatter plot of miRsift results for each miRNA family tested in multiple linear regression with pseudo bulk spermatogonia. miRsift uses single linear regression to test miRNA families for their contribution to RNA-seq changes individually and then multiple linear regression with significant to consider the effect of other miRNAs (https://github.com/SRHilz/miRsift). X-axis is regression coefficient from multiple linear regression test and y-axis is log10 FDR value for each miRNA. Negative regression coefficient represents decreased repression of miRNA family targets and positive regression coefficient represents increased repression of miRNA family targets. Red line indicates FDR significance threshold of 0.05. (F) As in (E) for leptotene/zygotene cluster. (G) As in (E) for pachytene

**Figure S10.**
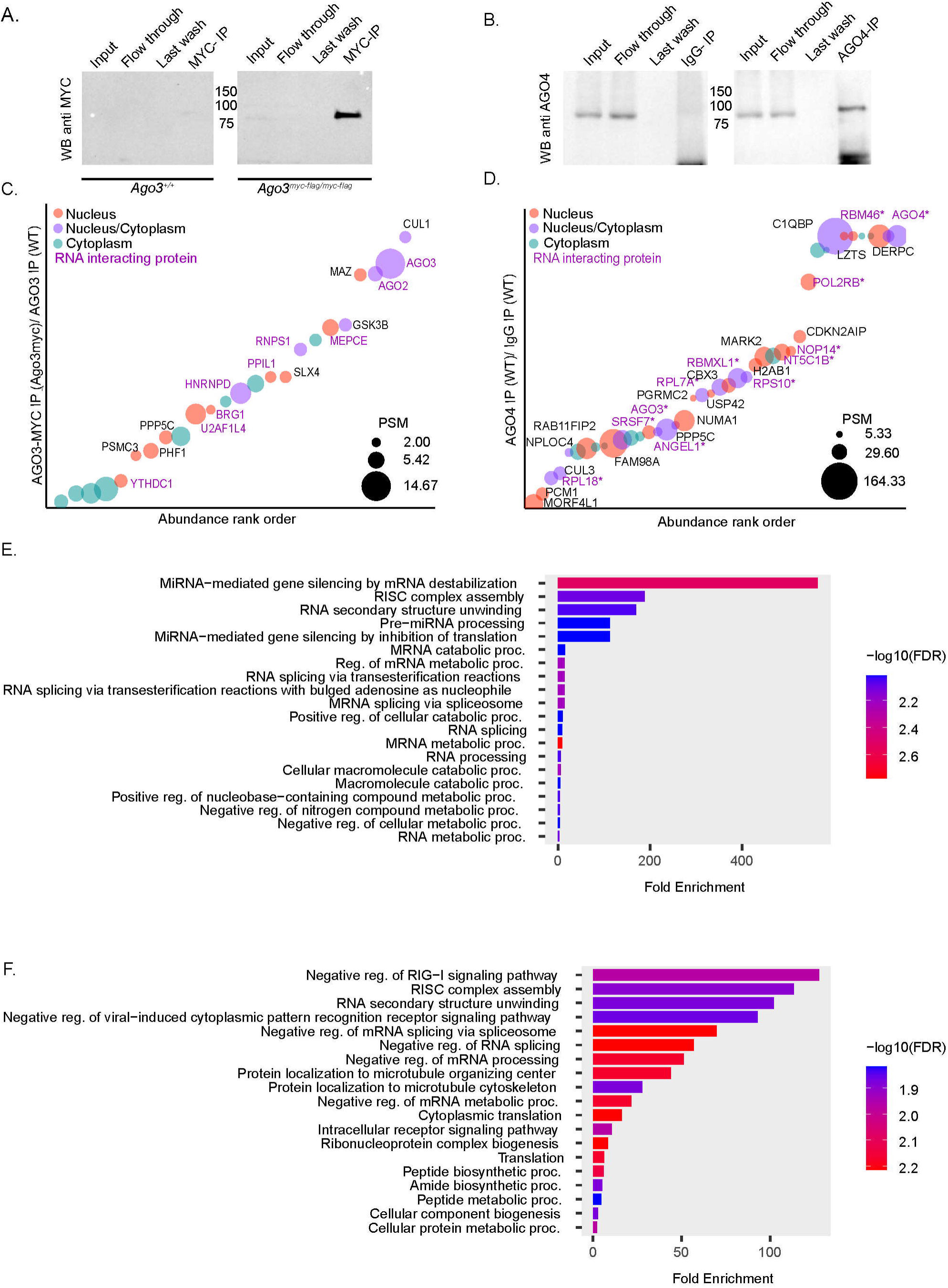
Potential AGO3 and AGO4 interactors in the mouse germline. (A) Immunoprecipitation of AGO3 using a MYC antibody in homozygous *Ago3^myc-flag^* germ cells and (B) AGO4 using an AGO4 antibody in wild type germ cells (n=3). (C) Bubble plot showing ranked abundance ratios for candidate protein interactors of AGO3 and (D) AGO4. For data visualization, we plot the log_10_-abundance ratios of each interactor against their abundance rank order. Every bubble represents an identified protein, and its size corresponds to the number of peptides found for that given protein. Abundance ratio was calculated as abundance of a given protein in the AGO IP relative to the negative control. (E) GO Analysis of candidate AGO3 and AGO4 (F) interactors.

**Figure S11.**
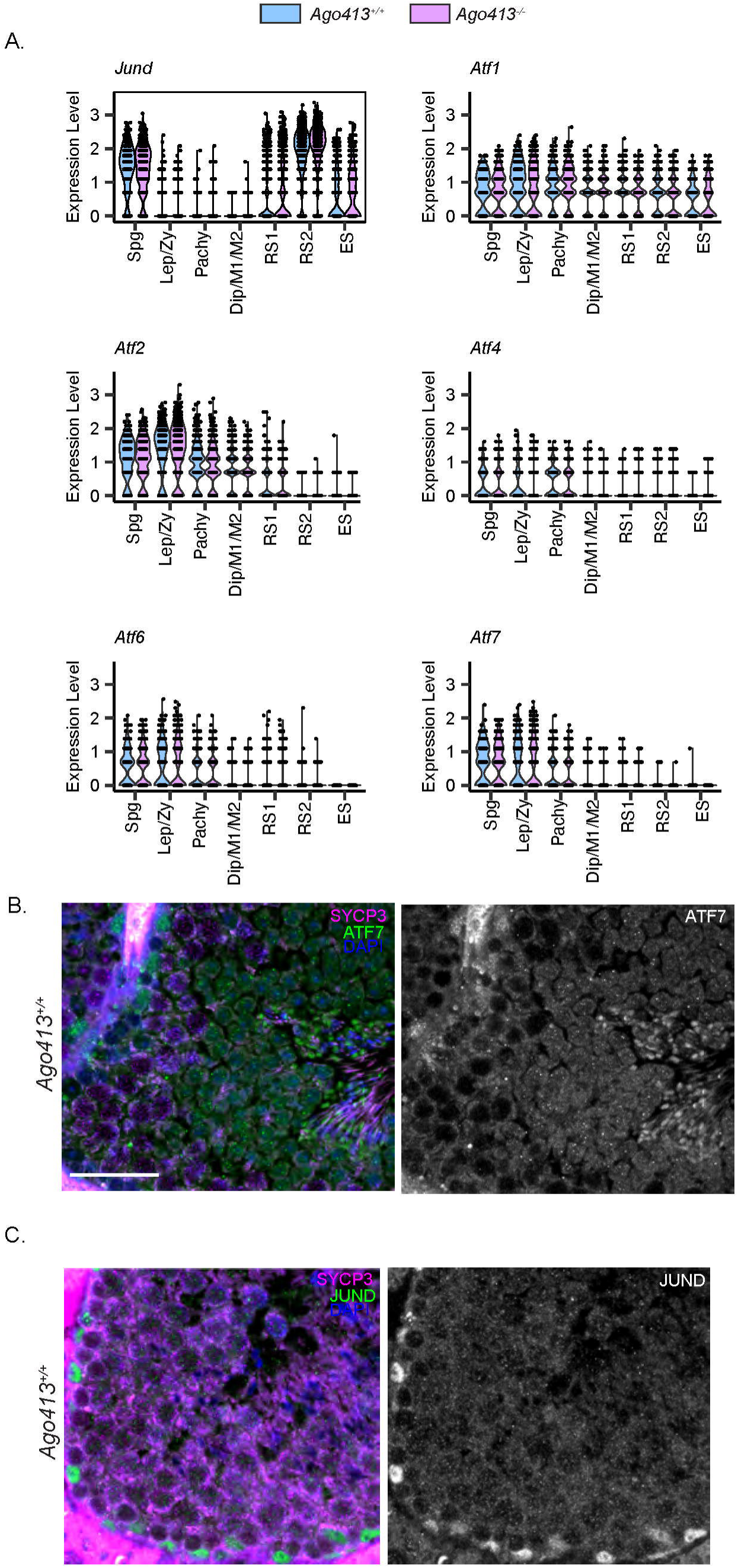
**(A)** Expression of AP-1 genes in *Ago413^+/+^* and *Ago413^-/-^* germ cell clusters. (B) Immunostaining of *Ago413^+/+^* testicular sections with ATF7 and SYCP3 (B) and JUND and SYCP3 (C). ATF7 shows diffuse localization to the spermatocyte and spermatid populations nucleus, while JUND is mostly enriched in the outer layer of the seminiferous tubules. Bars indicate 40 μm.

## METHODS

### Experimental animals

All mice used in this work were handled following institutional guidelines under the protocol 2013-0041, approved by the Institutional Animal Care and Use Committee (IACUC) at Cornell University. All mouse strains were backcrossed at least five generations and maintained on a C57BL/6J background (Jackson Laboratory). Mice were maintained under strictly controlled conditions of constant temperature, 12-hour light/dark cycles, and provided food and water *ad libitum*. All mice in this study were euthanized between 8 and 18 weeks by CO_2_ asphyxiation. Genotyping was conducted using ear snips obtained from mice at 4-6 weeks of age, followed by DNA extraction and PCR using the primers indicated on the resource table for each mouse line. The *Ago413* mouse line, strain #:014152, was obtained from Jackson Laboratory and maintained in our mice facilities by +/- by +/- crosses. This line was originally made in the 129 mouse strain and backcrossed into B6/C57 mice at least 4 times by the Hannon Lab, the donor lab. This lab validated the elimination of expression of all 3 Argonaute genes by quantitative RT-PCR of embryonic day 13.5 (E13.5) mouse embryonic fibroblasts. We also confirmed this by analysis of our germ cell RNAseq data, and Western Blot of protein lysates from testes of *Ago413^-/-^, Ago413^+/-^ and Ago413 ^+/+^* mice (Figure 1).

The *Ago3^myc-flag^* line was generated in the Cohen Lab by CRISPR/Cas9 knock in of the *myc/flag* coding sequence 41 base pairs upstream from the start codon, which corresponds to the N-terminal domain of the AGO3 protein. The single-stranded oligodeoxynucleotide used was the following: CGCTCCTCGCCTCTGTGGTGGCACCCTTCTCTCGTGAAGCACTCCCC (homology arm) <u>CCA (PAM)</u> GCTCCATGA (guide) ATG (start site) GAACAAAAACTTATTTCTGAAGAAGATCTG (myc) GACTACAAAGACGATGACGACAAG (flag) ATG (start site) GAAATCGG (guide) <u>CTCCGCAGGTGAGGCGAGCTGCGGGACAGGGCAGGTGGG (homology arm).</u>

The epitope tag was validated by Western blot, using protein lysates from whole testes and germ cells in *Ago3^my-flag/myc-flag^* and littermates *Ago3^+/+^* and an anti MYC tag antibody, where only *Ago3^myc-flag/myc-flag^* lysate showed a band running at ∼100 kDa (Figure 1).

### Immunoblotting

Protein extraction from whole testes or single germ cell suspensions was performed by lysing the cells in ice cold buffer (0.5M Tris-HCl pH 8.0; 1% NP-40; 150mM NaCl; 5mM EDTA plus protease inhibitors, Roche complete® tablet) followed by a sonication step (23 amplitude, 0.4 sec on, 0.2 sec off, 12 sec total). Then, the lysate was centrifuged for 20 min at 14.000g and the supernatant was recovered for protein concentration quantification using the Pierce BCA Protein Assay Kit (Thermofisher Scientific). For subcellular fractionation, the kit NE-PER (Thermofisher Scientific) reagent was used on single germ cell suspension following manufacturer’s protocol. Proteins were resolved by molecular weight in on homemade 12.5% bis-acrylamide gels or 4-20% MINIPROTEAN TGX pre casted gels (BioRad) and transferred to methanol activated PVDF membranes using a Biorad Mini Trans-Blot Celby. After transfer, membranes were blocked using EveryBlot Blocking Buffer (BioRad) for 10 minutes at room temperature and incubated with rocking overnight at 4°C with diluted primary antibodies in EveryBlot Blocking Buffer (see resource table for antibody information). After three consecutive washes in Tris-buffered saline with 0.1% Tween-20 (TBST) at room temperature for 5 minutes, membranes were incubated with secondary-HRP conjugated antibodies diluted in EveryBlot Blocking Buffer at room temperature for 2 hours. Finally, membranes were washed again three times in TBST and western blot signal was acquired with a Chemidoc Imaging System (BioRad).

### Prophase I chromosome spreads

Prophase I chromosome spreads were performed using the routine method employed by the Cohen Lab ^52,77,78^. Testes were detunicated, and tubules were placed for 20 minutes on ice on hypotonic buffer (30 mM tris; 50 mM sucrose; 17 mM sodium citrate; 5 mM EDTA; 5 mM PMSF; 2.5 mM DTT; pH 8.2-8.4). Then, tubules were minced in a 500 mM solution of sucrose at room temperature, and the single cell suspension obtained was spread onto a slide previously coated with 1% paraformaldehyde with 0.15% Triton X (pH 9.2-9.3). The slides were slowly dried for 2-3 hours in a humid chamber and let air-dry for an extra hour. Then, they were washed with 0.4% Photo-flo 200 solution (Kodak Professional) for 5 min. Staining was done by washing the slides in 0.4% Photo-flo in PBS for 10 minutes, followed by permeabilization for 10 minutes in 0.1% Triton X in PBS, and blocking for 10 minutes in 10% antibody dilution buffer (3% bovine serum albumin, 10% normal goat serum, 0.0125% Triton X, in PBS). Finally, slides were incubated overnight with primary antibodies diluted in antibody dilution buffer in a humid chamber at room temperature. Secondary antibody incubation was performed the next day with previous Photo-flo washing, permeabilization and blocking steps described for primary antibodies. Slides were incubated with secondary antibodies at 37°C in a humid chamber for 2 hours, washed in PBS with 0.4% Photo-flo three times plus one in 0.4% Photo-flo in distilled water and mounted in antifade media with DAPI (2.3% DABCO, 20 mM Tris pH 8.0, 8 μg DAPI in 90% glycerol). Slides were kept at 4°C until imaging on a Zeiss Axio Imager epifluorescence microscope equipped with Zeiss Zen Blue version 3.0 software (Carl Zeiss AG, Oberkochen, Germany). Images were processed using the Fiji software^79^. For fluorescence intensity quantification, cells were analyzed individually within their subpopulation. Background was subtracted of each image and a region of interest (ROI) was selected using Zen software to delimit each cell (whole nuclear fluorescence) and the sex body. Each image was saved individually and a FIJI macro script created in our lab was used as previously described ^52^to quantify the mean gray value (MGV) of intensity for each fluorescence channel corresponding to DAPI (DNA staining), BRG1 and SYCP3. To test for changes in mean gray value of BRG1, we used lme4 to fit a mixed model for log2 transformed gray value with sample as a random effect and cell type and genotype as a fixed effect with interaction (log2_intensity ∼ genotype*celltype + (1|sample)). We performed post-hoc testing of significant changes in estimated marginal means between cell types for each genotype with the emmeans package (emmeans (model, ∼ celltype|genotype)) and then did a pairwise comparison with Tukey test, using 95% confidence intervals.

### Immunofluorescence and apoptosis detection on testicular histological sections

To obtain histological sections of testes from each genotype, testes were removed after euthanasia, one of them fixed in 10% formalin and the other one in Bouins solution at room temperature for 6 hours followed by three sequential 5-minute washes in 70% ethanol. Fixed testes were later embedded in paraffin and processed to obtain histological sections, 5 µm thick. Histological sections were deparaffinized with Safeclear xylene substitutes (Fisher Scientific) followed by decreasing ethanol concentrations. The slides were then gradually dehydrated by incubation in increasing concentrations of ethanol.

For immunofluorescence, antigen retrieval was done by immersing the slides on boiling citrate buffer (10mM sodium citrate, 0.05% Tween 20, pH 6) for 20 min, followed by 20 min at room temperature. Then, slides were washed in PBS twice for 5 minutes. Sections were blocked in 0.05% Tween/PBS (PBST) with 1% bovine serum albumin and 3% goat serum for 1 hour at room temperature. After two washes with PBST for 5 minutes, sections were incubated with primary antibodies diluted in the blocking buffer at room temperature overnight. The next day, slides were washed three times in PBST and incubated with secondary antibodies diluted in blocking buffer for 2 hours at 37°C. Finally, sections were washed in PBST three times for 5 minutes and mounted in antifade media with DAPI (2.3% DABCO, 20 mM Tris pH 8.0, 8 μg DAPI in 90% glycerol).

Apoptosis levels in testes sections were evaluated with the commercial kit Apoptag kit (EMD Millipore) following the manufacturer’s instructions after deparaffinization. This kit detects DNA breaks associated with latest stages of apoptosis and is based on the dUTP nick-end labeling reaction of free 3’OH termini by terminal deoxynucleotidyl transferase (TdT), also known as TUNEL assay.

Testicular architecture was evaluated through Hematoxylin-eosin staining, followed by dehydration by sequential incubation in increasing concentrations of ethanol and mounting in toluene mounting media (Permount, Fisher Scientific). Hematoxylin-eosin stained and TUNEL sections were imaged using an Aperio CS2 Digital Pathology Slide Scanner microscope (Leica Biosystems) equipped with Aperio eSlide Manager Software. Immunofluorescent sections were imaged on a Zeiss Axio Imager epifluorescence microscope equipped with Zeiss Zen Blue version 3.0 software (Carl Zeiss AG, Oberkochen, Germany). Images were processed using the Fiji software ^79^.

### Testicular weight and sperm count

To study the reproductive phenotype in *Ago413* males, testicular weight and total body weight were recorded for each mouse after euthanasia. Both caudal epididymides were removed and placed in 1 mL of prewarmed and pH equilibrated DMEM containing 4% bovine serum albumin (Sigma-Aldrich). Sperm was allowed to swim out of the epididymis by incubation for 20 min at 37°C under 5% CO_2_ atmosphere. After this period, an aliquot of sperm was obtained and fixed in 10% formalin for quantification of total sperm using a hemocytometer.

### Enrichment on germ cells by Fluorescence activated cell sorting (FACS)

To prepare enriched germ cells fractions from each genotype for scRNA seq, we used Fluorescence activated cell sorting following the method developed by Rodriguez-Casuriaga et al ^80^. First, a single cell suspension from testes was obtained following the method described by Ascenção et al ^33^. Testes from 2-5 mice from the same genotype were collected, the *tunica albuginea* was removed, and tubules were placed in 10 ml of preheated (37°C) DMEM-F12 containing 2 mg of Collagenase 1A (Sigma) for 2 minutes with manual shaking. Digestion was stopped by two washes of the tubules with DMEM-F12. Next, the tubules were incubated in 10 ml of DMEM-F12 with 5 mg of trypsin (Sigma) and 7 mg/ml of DNAse I (Sigma) at 37°C with constant shaking (150 rpm) until the length of the tubules was about 1 mm (approximately 10 minutes). Digestion was stopped by addition of 3 ml of Fetal Bovine Serum (FBS, Sigma) and 30 ml of DMEM F-12. Digested tubules were strained on a 100 μm strainer and centrifuged at 15°C for 5 minutes at 600g. The single cell suspension of germ cells obtained was then resuspended in FBS and ^7931^incubated with 10 μM of Vybrant dye cycle (VDG) (Invitrogen) for 30 min at 37°C in the dark with constant rocking. VDG is DNA-specific vital stain that is fluorescent upon binding to double-stranded DNA and is excited at 488 nm with emission ∼520 nm. Sorting of cells was performed at Cornell Flow Cytometer Facility (RRID:SCR_021740), using a Sony MA900 fluorescent activated cell sorter, tuned to emit at 488 nm and laser power set to 100 mW to collect for VDG positive cells. Flow cytometric profiles were obtained by representing Forward Scatter (Y axes) vs VDG fluorescence intensity (X axes), this allowed to identify different cell populations in the sample based on their DNA content and size. To enrich for spermatocytes for single cell RNA-seq, a sorting gate was set to collect all germ cell types excluding sperm ^80^. Sorted cells were recovered in tubes containing DMEM-F12 buffer and 10% FBS. Enrichment in germ cells of the sorted cell populations was evaluated by performing immunofluorescence of meiotic spreads with the markers SYCP3, γH2AX and DAPI as previously described.

### Single Cell RNA sequencing library preparation

Single cell RNA sequencing libraries were prepared from enriched germ cell suspensions at the Transcriptional Regulation and Expression Facility at Cornell (RRID:SCR_022532) using the 10X Genomics Chromium Single Cell 3′ RNA-seq v3.1 kit. Flow sorted cells were processed on the 10X Genomics Chromium X System, targeting a total of 7000 cells per sample. Quality control was evaluated using an Agilent Fragment Analyzer and ran on a NovaSeqX platform with 150 base-pair reads.

### Single-cell transcriptome analysis

Fastq files were run through cellranger count (10x Genomics) [cellranger count –id=sampleID -- r1-length=28 --transcriptome=/path/to/refdata-gex-mm10-2020-A --fastqs=/path/to/directory -- sample=file_prefix --localcores=8 --localmem=64] to generate count tables. Ambient RNA correction was then performed on samples using cellbender [cellbender remove-background -- input /path/to/raw_feature_bc_matrix.h5 --output /path/to/output.hd5 --projected-ambient-count-threshold 0 --cpu-threads 32 --fpr 0.0 0.01 0.05 0.1 0.2 0.3 (--learning-rate 0.000025 for WT3; --learning-rate 0.00005 for KO1)]. Corrected samples with fpr 0.01 were imported into R to analyze with Seurat using Read_CellBender_h5_Multi_File (scCustomize). Cells were filtered based on number of genes detected, number of UMIs, and mitochondrial percentage, and cells with less than 500 genes or more than 9000 genes, less than 1500 UMIs, or more than 5% mitochondrial reads were removed from the analysis. Cell cycle scoring was performed (CellCycleScoring using Seurat genes and default parameters) and was used to calculate the difference between the S phase and G2M phase scores. Cell counts were normalized (SCTransform, vars.to.regress=“CC.Difference”, return.only.var.genes=F) and used to run PCA analysis (default parameters). Batch correction was performed using RunHarmony and cells were then clustered (FindNeighbors; dims = 1:12, reduction = “harmony”, k.param = 30; FindClusters; resolution = 0.3, n.start = 100). Cluster cell types were manually identified using expression of known marker genes (*Cenpa*, *Stra8*, and *Dazl* for spermatogonia; *Dazl*, *Gm960*, and *Meiob* for leptotene and zygotene spermatocytes; *Psma8* and *Piwil* for pachytene; *Pou5f2* and *Ccna1* for diplotene and dividing spermatocytes; *Tex36*, *Sun5*, *Tssk1*, *Cstl1*, *Tnp1*, and *Prm1* for different stages of round spermatids; *Tnp1*, *Prm1*, lack of *Tssk1*, and lower *Cstl1* for elongating spermatids; and *Col1a2*, *Acta2*, *Vcam1*, *Insl3*, *Laptm5*, *Hbb-bt*, *Ptgds*, and *Wt1* for somatic cell types).

Doublets were then removed using DoubletFinder. To remove doublets, counts were normalized for each sample using SCTransform (default parameters) and then a PCA (default parameters) and UMAP (dims=1:10) analysis were run. pK value for doublet removal was then chosen using paramSweep (PCs=1:10, sct = TRUE), summarizeSweep, and find.pK (default parameters), with chosen pK of for WT1, for KO1, for WT2, for KO2, for WT3, and for KO3. Homotypic doublets were modeled using modelHomotypic with the previously identified cell types and doublet removal was then run (doubletFinder using parameters PCs = 1:10, pN = 0.25, pK = chosen pK, nExp = round(0.045*#cells), reuse.pANN = FALSE, sct = TRUE and then rerun reusing the pANN and nExp = round(round(0.045*#cells)*(1-homotypic proportion)) with other parameters the same). Once doublets were removed, the samples were renormalized using the same parameters, PCA was run (default parameters), and samples were integrated using Harmony (IntegrateLayers; method = HarmonyIntegration). Samples were clustered (FindNeighbors; dims = 1:25, reduction = “harmony”, k.param = 30; FindClusters; resolution = 0.3, n.start = 100). Cluster cell types were identified as previously described. One cluster showed very low expression of marker genes and expression of multiple marker genes which are not expressed in the same stage of cells and were removed. Normalization, PCA and integration were rerun on the remaining cells with the same parameters. Samples were reclustered (FindNeighbors; dims = 1:15, reduction = “harmony”, k.param = 30; FindClusters; resolution = 0.4, n.start = 100), for a total of 11 clusters, the UMAP projection was created (RunUMAP; dims = 1:15, reduction = “harmony”, n.neighbors = 30), and cluster cell types were identified and used in all following analysis. Cell type proportion changes were tested using sccomp ^81^ (sccomp_glm with parameters formula_composition = ∼genotype and bimodal_mean_variability_association = TRUE). G2/M cell cycle correlations were produced by taking either the subset of cells in a cluster or all cells and using a Spearman correlation to between the G2/M correlation and the SCTransform normalized counts for each gene in the G2/M cell cycle list. SCTransform normalized counts were used for mean expression levels by cell. Both values were visualized using ComplexHeatmap (parameters row_km = 5, row_km_repeats = 100). Chromosome X and chromosome 9 ratios were calculated for each cell by taking the mean SCTransform normalized expression of all genes on either chromosome X or chromosome 9 and dividing by the mean expression of all autosomal genes. Ratios were visualized using ggplot and the gghalves package (geom_half_violin).

### WT cell type marker genes

Marker genes were identified for each cell type in wildtype cells (PrepSCTFindMarkers; FindAllMarkers, test.use = “MAST”, recorrect.umi = F). Marker gene sets for enrichment were then constructed by subsetting marker genes for adjusted p-value < 0.01 and average log2 fold-change >= 2 and then taking the top hundred genes ranked by adjusted p-value. Any genes that were duplicated between cell types were removed from the lists.

### Pseudobulk analysis

Cellbender corrected counts were aggregated by sample and cell type, as well as by cluster for cells identified as round spermatid 1 and 2. Counts were separated by cell type and filtered to remove genes with less than three samples having 10 or more counts. DESeq2 was used to normalize counts and to perform differential expression testing between knockout and wildtype samples with batch included in the design matrix for all samples and cluster also included for round spermatid 1 and 2 tests. The log_2_ fold-change results were used to run gene set enrichment analysis using clusterProfiler for GO terms (gseGO; ont= “BP”, OrgDb = org.Mm.eg.db, minGSSize = 50, pvalueCutoff = 0.05, verbose = FALSE, keyType = “ALIAS”) as well as for the marker genes identified for different cell types in wild type cells (GSEA; minGSSize = 20, pvalueCutoff = 1, verbose = FALSE). All genes with adjusted p-value <0.05 and absolute value log_2_ fold-change > 0.5 were considered significantly differentially expressed for the DE gene barplots.

### Bulk RNA sequencing and small RNA sequencing

Bulk RNA sequencing and small RNA sequencing was performed in enriched fractions of pachytene spermatocytes and round spermatids obtained using a BSA gradient method developed by Da Ros and others ^34^. Briefly, a single germ cell suspension was prepared from testes of each genotype following the published protocol and loaded into a 5-layer discontinuous bovine serum albumin density gradient for 2.5 hours to allow enrichment of the different cell populations by gravity sedimentation. The cell fractions were then manually collected, and cells were pelleted and washed in cold PBS by centrifugation. Purity of each fraction was evaluated by light microscopy and confirmed by immunofluorescence of meiotic spreads with the markers SYCP3 γH2AX and DAPI as previously described, and only fractions containing 70% to 80% purity for pachytene/diplotene spermatocytes and 80 to 90% purity for round spermatids were pooled. Finally, cells were resuspended in TRIzol LS (Thermo Fisher Scientific) and snap frozen for downstream RNA extraction with an extra chloroform extraction to remove residual phenol and addition of glyco-blue as a carrier to promote RNA precipitation. RNA integrity was assessed on an Agilent Fragment Analyzer.

Directional RNA-seq libraries were prepared from 100-200ng total RNA by the Transcriptional Regulation & Expression Facility at Cornell using the NEBNext Directional Ultra II RNA Library Prep Kit for Illumina (New England Biolabs), with initial polyA+ selection. Library quality was assessed on an Agilent Fragment Analyzer and libraries were sequenced on an Illumina Novaseq 6000 with 2×150bp reads.

Small RNA-seq libraries were prepared from 100ng total RNA by the Transcriptional Regulation & Expression Facility at Cornell using the NEBNext Small RNA Library Prep Kit for Illumina (New England Biolabs). Library quality was assessed on an Agilent Fragment Analyzer and libraries were sequenced on an Illumina NextSeq 500with 50 bp single end reads.

### Bulk RNA-seq analysis

Illumina pipeline software was used for base calling. Sequenced reads were trimmed for 3’ adaptor sequence and low-quality sequence and filtered to remove reads < 50nt with TrimGalore. Processed reads were mapped to the reference genome with STAR using --quantMode GeneCounts to generate raw counts per gene. Raw counts were then imported into R and subset by cell type for differential expression analysis. Any genes without at least 10 counts in at least 3 samples were pre-filtered and DESeq2 was used to normalize the data and compare expression between knockout and wild-type samples. The log_2_ fold-change values were used for gene set enrichment analysis using clusterProfiler with the wild-type scRNA-seq marker gene sets (GSEA; minGSSize = 20, pvalueCutoff = 1, verbose = FALSE). Genes with adjusted p-value <0.05 and absolute log_2_ fold-change > 0.5 were considered significantly differentially expressed for DE bar plots.

### smRNA-seq analysis

Illumina pipeline software was used for basecalling. Sequenced reads were trimmed for 3’ adaptor sequence and low-quality sequence and filtered to remove reads < 10nt with TrimGalore. Trimmed reads were processed using from miRDeep2 pipeline: collapsed to fasta format using fastq2fasta.pl, processed using mapper.pl, and mapped to miRBase^82^ v22.1 mmu miRNA using quantifier.pl. Raw counts were imported into R and counts from individual miRNA were summed into families as defined by TargetScan Mouse v8 ^83^. Counts tables were subset by cell type and miRNA families were removed that did not have at least 10 counts in at least 3 samples. DESeq2 was used to normalize the data and compare between knockout and wildtype samples. Log_2_ fold-changes and expression values were used in post-transcriptional regulatory analysis.

### Sperm motility and morphology

Sperm Motility was assessed using computer aided sperm analysis (CASA). An aliquot of sperm obtained by swim-out was collected as described above using TYH media ^84^ without sodium bicarbonate and or BSA and placed on a cytometry chamber mounted on a warmed stage (37°C). Sperm motility was recorded using 40X magnification and the percentage of motile sperm over total sperm per sample was calculated using software Hamilton Thorne HT CASA SCA. At least five different fields and 200 sperm cells were recorded. Sperm morphology was assessed in smears of fixed sperm with 1% paraformaldehyde and stained with hematoxylin and eosin ^85^. A minimum of 250 sperm were visualized per mouse.

### Breeding assay, blastocyst collection and sexing of progeny

Homozygous wild type, heterozygous and homozygous null *Ago413* males of 10 weeks old were housed to breed to CD-1 females. After natural mating, time to first litter, litter size and sex ratio were recorded. The breeding pair was kept together until they produced at least a total of three litters. For blastocyst collection, breeding pairs of *Ago413* males from the three genotypes and CD-1 females were housed together and females were checked for plugs every morning. Upon plugging, females were removed from the breeding and housed individually until 3.5 days post plug. At this point, females were euthanized, and the uterus was quickly dissected out and placed directly in a watch glass containing pre-warmed M2 media (MR-015-D, Sigma-Aldrich) in a heated dissection hood. Blastocysts were flushed from the uterus by inserting a 1ml syringe filled with M2 media into the lumen of the uterus just below the oviduct and pushing the fluid through the uterine horns and out the cervix. Blastocysts were collected from the watch glass, washed in a droplet of PBS and individually pipetted into PCR strip tubes and frozen at -80 °C. After thawing, DNA was extracted from blastocysts by using the Extract-N-Amp Tissue PCR Kit (XNAT2, Sigma-Aldrich). Female and male embryos were identified by PCR using primers designed flanking an 84 bp deletion of the X-linked *Rbm31x* gene relative to its Y-linked homolog *Rbm31y* as previously described ^86^.

### Digital Droplet PCR (ddPCR) of sperm DNA

Sperm DNA extraction was adapted from Cole and Jasin ^87^. Spermatozoa was prepared using the swim up technique that allows to separate motile from non-motile spermatozoa based on its ability to swim upwards through an overlaid medium ^88^. Swim up and non-swim up sperm were passed through a 70 µm filter washed four times with saline sodium citrate by centrifugation at 5000 rpm for 1 minute. Sperm was resuspended in 5% SDS, 50% β-mercaptoethanol, 0.002% of proteinase K, in sodium citrate buffer and incubated for 2 hours at 55 °C inverting occasionally. Following incubation, a volume of phenol: chloroform: isoamyl alcohol (25:24:1) was added to extract protein, and tubes were spun at 15000 xrcf for 5 minutes. The aqueous phase was then transferred into a new tube and DNA was precipitated by adding ice cold ethanol. The sample was mixed well and spun at 15000 x rcf for 5 minutes, the supernatant was discarded, and the pellet was washed in 70% ethanol by repeating centrifugation and removing the supernatant with a pipette tip. The pellet was then resuspended in DNAse free water and washed in 1/10 volume of sodium acetate and 3 volumes of cold ethanol and centrifuged again. The pellet was then washed once again in 70% ethanol, centrifuged and the supernatant discarded. The pellet was then left to air dry before being suspended in 100uL 5mM Tris HCl and incubated for 1 hour at 55°C. Samples were stored at -20°C until digital droplet PCR (ddPCR). To target the Y and X chromosome, TaqMan Gene Expression Assay probes (Thermo Fisher Scientific) against the genes *Sry* with a FAM label and *Rbmx* with a VIC label were used respectively. ddPCR was carried out by the Genomics Facility at Cornell (RRID:SCR_021727). The ratio of VIC to FAM signal was calculated to give the ratio of X to Y bearing sperm present in each sample.

### DNA Fluorescence in-situ hybridization (FISH) on sperm

Sexing of sperm was performed by targeting the X and Y chromosome for Fluorescent in situ hybridization (FISH) using an adapted protocol from Whyte et al ^89^. Sperm was collected by swim up procedure as previously described and fixed in pre-soaked slides in ethanol for 48 hs. Slides were then dehydrated in 80% methanol at -20 °C for 20 minutes and then again air-dried overnight at room temperature. A droplet of 10mM dithiothreitol (DTT) in 0.1M Tris-HCl was added to the slides and then microwaved for 15 seconds at 550 Watts. Next, slides were drained and 20 μL of 10mM lithium diiodosalicylate (LIS) and 1mM DTT diluted in 0.1 mM Tris-HCl was added to each well. Slides were then covered in parafilm, microwaved for 90 seconds at 550 Watts and then washed twice in 2X saline sodium citrate. Slides were then treated again with 80% methanol for 20 minutes at -20 °C and air dried at room temperature. X and Y chromosome probes (Empire Genomics MCEN-XY-10-GRRE) were added to the hybridization buffer (Empire Genomics) and denatured at 65°C for 10 mins and then held at 37 °C for 30 minutes. Probes were then added to each well, coverslips were applied and sealed with rubber cement and then microwaved for 78 seconds at 1100 Watts. Slides were transferred to a 37 °C prewarmed humid chamber and left for 18 hours at 37 °C to complete hybridization. Following hybridization, slides were washed in a pre-warmed 45 °C citrate buffer for 5 minutes to loosen coverslips. Slides were then placed into a pre-warmed wash solution of 50% formamide with 50% sodium citrate and kept at 45 °C for 8 minutes, drained and mounted with DAPI antifade. Slides were imaged on a Zeiss Axio Imager epifluorescence microscope equipped with Zeiss Zen Blue version 3.0 software (Carl Zeiss AG, Oberkochen, Germany). At least 200 sperm cells were counted and classified as X- or Y-bearing sperm.

### *In-vitro* fertilization (IVF)

Three-week-old C57BL/6J wild type females were super ovulated by intraperitoneal injection of 5IU pregnant mare serum gonadotropin (PMSG, Sigma), followed by 5IU of human chorionic gonadotropin (HCG, Sigma Aldrich) 46 hours later. Ovulated females were humanely euthanized, and ovaries were dissected out and placed in a watch glass containing pre-warmed M2 media (Milipore-Sigma, MR-015). *Cumulus*-oocyte clusters (COCs) were removed from each ovary and placed in pre-warmed M2 media and dissociated using hyaluronidase. Sperm was obtained by swim out in pre-equilibrated HTF media (Milipore-Sigma, MR-070-D) on a 5% CO_2_, 5% O_2_ and 90% N_2_ incubator at 37 °C for 30 minutes. After this time, sperm were counted using a hemocytometer and an equal number of sperm from each male was added to a 250μl droplet of HTF to achieve a concentration of 2 x 10^6^ spermatozoa/ml. Dissociated COCs were then transferred along with minimal media to the droplet containing sperm and incubated at 37 °, 5% CO_2_, 5% O_2_ and 90% N_2_ for 4 hours. After this time, COCs were transferred to plates containing warmed pre-equilibrated KSOM media (Millipore-Sigma, MR-101) and analyzed for the presence of a second polar body.

### Post-transcriptional regulatory analysis

Spermatocyte and round spermatid miRNA sets for the targeting signature analysis were chosen by taking the top 25 expressed miRNA and the top 10 expressed miRNA that were differentially expressed from each bulk smRNA-seq differential expression analysis. For spermatogonia, leptotene/zygotene, pachytene, and diplotene/dividing cell types the spermatocyte miRNA set was used, and four round spermatids 1 and 2 and elongating spermatids the spermatid miRNA set was used. The targeting signature was tested using the log_2_ fold-change from the pseudobulk analysis by subtracting the mean log_2_ fold-change of targets (TargetscanMouse v8 total weighted context score for gene < -0.2) from the mean log_2_ fold-change of background genes (genes without any predicted target sites for miRNA family but at least one predicted target site for any other miRNA family). Targeting signature significance was tested using a Wilcoxon rank sum test. The same process was used for spermatocyte and round spermatid bulk RNA-seq testing. For spermatid development gene set and XY-linked gene targeting analysis log_2_ fold-change of miRNA targets as defined above were plotted with a background set of all genes in the spermatid development gene set without a target site for any of the miRNA families plotted in the CDF plot.

To run miRsift, first buildContextTable was run with modified code to include all miRNA families which are broadly conserved and all miRNA families which are expressed in the smRNA-seq data and Targetscan identifies as conserved or poorly conserved but confidently annotated. For each of the cell types, pseudobulk counts were imported and prepared for analysis using importSeq and then the linear regression was run using miRsiftAnalyze.

### Immunoprecipitation and Mass Spectrometry (IP-MS)

Immunoprecipitation was done in germ cell enriched single cell suspensions obtained from mouse testes as previously described^77^. After removing the testicular *tunica albuginea*, the seminiferous tubules were minced in PBS and further dissociated by pipetting to release the germ cells. The tubules and cell suspension were then filtered through a 70 μm cell strainer and the cells were pelleted by centrifugation at 4°C for 5 minutes at 600g. The cell pellet was resuspended in protein lysis buffer (50 mM Tris, 0.2% NP-40, 150 mM NaCl, 5 mM EDTA) with EDTA-free Protease Inhibitor Cocktail (Sigma) and sonicated for 12 seconds at 23% amplitude in cycles of 0.4 seconds on and 0.2 seconds off. Finally, lysates were centrifuged at 4 °C for 20 minutes at 15,000 x g, and the supernatant collected in a new tube. Before immunoprecipitation, lysates were precleared by incubation with the corresponding magnetic beads with no antibody conjugated (see resource table) for 1 hour at 4°C on a nutator. After this period, beads were pelleted using a magnet and the precleared lysate was transferred to a new tube. For AGO4 immunoprecipitation, a lysate made from C57/B6 germ cells was incubated with an anti AGO4 antibody overnight at 4 °C on a nutator and an anti IgG rabbit was used as a control. The next day, the lysate was incubated with 10 μg of Protein A Dynabeads (Invitrogen) for 2 hours at 4 °C on a nutator. For AGO3 immunoprecipitation, a lysate made from *Ago3^myc-flag/myc-flag^* germ cells was incubated with 10 μg of anti-MYC antibody covalently bound to magnetic beads (Cell Signaling) overnight at 4 °C on a nutator. As a control, lysates from *Ago3^+/+^* germ cells were incubated with the same concentration of MYC-beads. After antibody and bead incubations, the flow through was collected and beads were washed six times with cold lysis buffer by pipetting, followed by pelleting using a magnet. Following the final wash, beads were resuspended in elution buffer (100mM Tris, 1% SDS, 10mM DTT) and incubated at 65 °C for 15 minutes. After incubation, the beads were pelleted with a magnet and elution was collected and used for Western Blot and Mass Spectrometry Analysis and label-free quantitation (LFQ) by the Proteomics and Metabolomics Facility at Cornell (RRID:SCR_021743).

Elutions of three replicates per each immunoprecipitation were analyzed by nano LC-MS/MS using an Orbitrap Fusion mass spectrometer (Thermo-Fisher Scientific, San Jose, CA) equipped with a nanospray Flex Ion Source using high energy collision dissociation (HCD) coupled with UltiMate3000 RSLCnano (Dionex, Sunnyvale, CA). Reconstituted samples were injected onto a PepMap C-18 RP nano trap column (3 μm, 100 μm x 20 μm, Dionex) with nanoViper fittings at a 20 μL/min flow rate for on-line desalting, and subsequently separated on a PepMap C-18 RP nano column (3 μm, 75 μm x 25 μm) and eluted in a 120 min gradient of 5-35% acetonitrile (CAN_ in 0.1% formic acid) at 200 nL/min. The Orbitrap Fusion was operated in a data-dependent acquisition mode using FT mass analyzer for one survey scan followed by three “Top Speed” data-dependent CID ion trap MS/MS scans with normalized collision energy of 30%. A dynamic exclusion window of 45 seconds was specified. Data were acquired using Xcalibur 3.0 operation software and Orbitrap Fusion 2.0 (Thermo-Fisher Scientific).

Raw spectra for each sample and replicate were processed and proteome databases searched using Proteome Discoverer 2.5 (PD 2.5, Thermo-Fisher Scientific) with the Sequest HT search engine. Identified peptides were filtered for a maximum of 1% false discovery rate (FDR) using the Percolator algorithm in PD2.5. Relative label-free quantification was done in PD 2.5 to calculate protein abundances. The number of peptide spectrum matches (PSMs) were summed to represent protein abundance. Abundance ratios were calculated based on pairwise ratios using the median calculated among replicates.

### Motif Analysis

To identify motifs of potential transcription factors associated with gene expression changes, transcriptional regulatory regions previously identified using dREG (cite here) analysis of leChRO-seq data ^52^ were used. All peaks within 2kb of genes with adjusted p-value < 0.05 and log_2_ fold-change >= 0.05 in the pseudobulk differential expression analysis were used for motif analysis with Homer using the 100 bp region on either side of the peak summit (findMotifsGenome.pl peaks.bed mm10 output_dir/ -size 200 -len 8,10,12 -p 4). Peaks were also subset to only include those within 2kb of significantly upregulated genes which were on the X and Y chromosomes, and motif analysis was run with the same parameters.

